# Expansion of Ventral Foregut Primes the Enhancer Landscape for Organ Specific Differentiation

**DOI:** 10.1101/2022.04.11.487673

**Authors:** Yan Fung Wong, Yatendra Kumar, Martin Proks, Jose Alejandro Romero Herrera, Michaela Mrugala Rothová, Rita S. Monteiro, Sara Pozzi, Rachel E. Jennings, Neil A. Hanley, Wendy A. Bickmore, Joshua M. Brickman

## Abstract

Cell proliferation is fundamental for almost all stages of development and differentiation that require an increase cell number. Although cell cycle phase has been associated with differentiation, the actual process of proliferation is not seen as having a specific role. Here we exploit human embryonic stem cell derived endodermal progenitors that we find are an *in vitro* model for the ventral foregut. These cells exhibit expansion dependent increases in differentiation efficiency to pancreatic progenitors that are linked to organ-specific enhancer priming at the level of chromatin accessibility and the decommissioning of lineage inappropriate enhancers. Our findings suggest that cell proliferation in embryonic development is about more than tissue expansion, it is required to ensure equilibration of gene regulatory networks allowing cells to become primed for future differentiation. The use of expansion of lineage specific intermediates may therefore be an important step in high fidelity *in vitro* differentiation.

## Introduction

The regulation of gene expression during differentiation is generally thought of as a linear process that involves the action of extra-cellular signalling and transcription factors (TFs) on gene expression. Cell proliferation is regarded as peripheral to differentiation itself, although it has a clear function in the selection of specific cell types. While cell cycle phase has been linked to lineage specific differentiation^1, 2^, here we explore the notion that differentiation requires proliferation to enhance the processing of lineage promoting information.

As there are a limited number of conserved signal transduction pathways but a large diversity of developmental outcomes from these pathways, the means by which cells regulate their competence to differentiate and manifest lineage specific outcomes is determined by the TF code expressed in these cells and the activity of specific TFs bound regulatory elements known as enhancers. While there is a large body of data on the signalling and TF networks during directed differentiation of pluripotent stem cells to specific lineages^3, 4^, here we focus on how progenitor cell states stabilize transcriptional priming.

In embryonic development the visceral organs are formed from a progenitor population known as the endoderm^5^. These cells are initially specified during gastrulation and undergo extensive proliferation as they prepare to differentiate into distinct organ primordial^6^. In particular, the liver and pancreas are derived from the anterior definitive endoderm (ADE). ADE is formed as a result of the anterior migration of cells from the anterior region of the primitive streak at the beginning of gastrulation. The anterior-most DE will then migrate ventrally to form the ventral foregut, containing a bipotent precursor of liver and ventral pancreas^7, 8^, a population that has recently been shown to expand and retain potency for both lineages *in vivo* over a period of several days of mouse development^9^.

Embryonic stem cell (ESCs) are pluripotent stem cells derived from the pre-implantation embryo that can be differentiated *in vitro* to form all embryonic germ layers including endoderm^10, 11^. The directed and linear differentiation of ESCs has been used to produced organ-specific cell types such as pancreatic beta cells^12–14^ and hepatocytes^15, 16^. In addition, the generation of expandable endodermal progenitors (EPs) derived from both mouse and human ESCs (hESCs) can be used as a staging platform for further differentiation^17–19^.

The expansion of endodermal cells from hESCs has been shown to specifically promote the generation of more mature pancreatic beta cells^17^. Here we characterise these expanding EPs and find that they are the *in vitro* equivalent of the ventral foregut, suggesting they represent a human cell culture model for this region of the embryonic gut. Consistent with their developmental identity, they can readily be converted into expanding pancreatic spheroids or hepatic organoids, and can be further differentiated with a high efficiency that is linked to expansion. We show that expansion or continued proliferation results in organ-specific lineage priming that is correlated to levels of organ-specific enhancer accessibility. Enhancer priming is not accompanied by large changes in transcription of organ-specific genes, but instead prepares these enhancers for their activation in response to signalling. Expansion of ventral foregut culture also appears to ensure that enhancers for lineages outside of those normally produced by the ventral foregut are decommissioned. The net result is that further differentiation allows lineage appropriate enhancers to engage transcriptionally and their enhanced state of readiness was found to be regulated by the presence of sequence specific transcription factors that act on these elements over the course of multiple cell cycles. Our findings suggest that the extensive cell proliferation that characterises normal embryonic development is not merely required for tissue expansion, but to ensure equilibration of gene regulatory networks allowing cells types to become primed for future high-fidelity differentiation.

## Results

### Expanding endoderm progenitors as an *in vitro* model for ventral foregut

To characterise the impact of expansion on the differentiation competence of human endodermal cell types, we focused on three dimensional EP culture^17^ derived from two independent hESC lines, H9 and HUES4. This protocol expands endoderm in the presence of FGF2, BMP4, VEGF, EGF^17, 19^, cytokines known to act in the ventral foregut region. We quantitated gene expression during expansion by single cell RNA-seq and showed that transient ADE cells comprise two sub-populations (ADE.1 and ADE.2) while EP culture was homogenous (Extended Fig. 1a, left). In human development, formation of ventral foregut endoderm has been detected at Carnegie Stages 8-9^20^. To compare populations in these distinct datasets, we used our Cluster Alignment Tool (CAT) (Rothova et al, submitted) to compare gene expression changes in EP cells to expression within regions of human embryonic gut from these stages: human foregut (hFG.1-4), the lip formed from ventral foregut, referred to as the lip of the anterior intestinal portal (hAL), midgut (hMG.1-3), and hindgut (hHG.1-2)^21^ at single-cell level (Extended Fig. 1a, right). We found that ADE clusters align to the foregut hFG.2 cluster and midgut hMG1 cluster (Fig. 1a). In contrast, EP cells align with the hAL cluster in addition to the midgut cluster picked by ADE, hMG1, that represents a population cells located immediately adjacent to the hAL^21^ (Fig. 1a). As EP cells align to both clusters, we assessed the expression of genes that are specifically enriched in RNA-seq dataset of H9 EP cells (Extended Fig. 1b), and then asked if this set contained genes that were differentially expressed when the hAL and hMG.1 clusters were compared, with a few exceptions, genes expressed at higher levels in hAL, were also elevated in EP cells (Fig. 1b). To further confirm the hAL or ventral foregut identity of EP cells, we observed the expression of the hAL markers HHEX^21^ and TBX3^17^ by immunohistochemistry (Extended Fig. 1c).

**Fig. 1:**
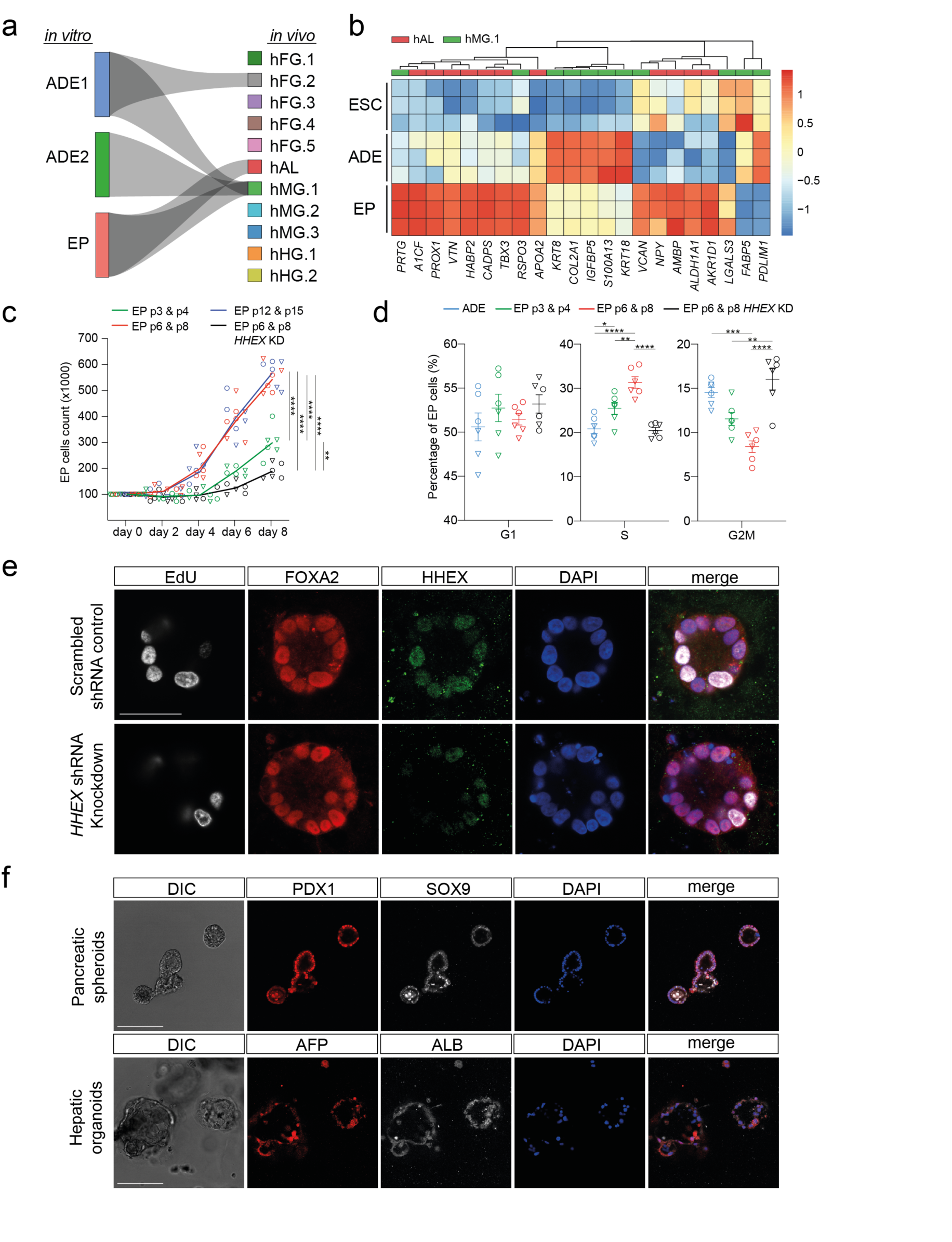
Expanding endoderm progenitors as an *in vitro* model for ventral foregut. **a**, Sankey diagram visualizing the CAT alignments between *in vitro* clusters (ADE.1, ADE.2, and EP) from this study and *in vivo* endodermal clusters from the Li et al. dataset^21^. Only significant CAT alignments between clusters are shown. **b**, Heatmap showing expression of hAL and hMG marker genes in ESC, ADE and EP cells (Bulk RNA-seq dataset, log normalised counts, N = 3). Only markers expressed significantly different between ADE and EP are shown (log2 fold change > 1.5, adjusted p-value < 0.05). **c**, Cumulative growth curves showing EP cell counts at different passages of expansion for control and HHEX knockdown (EPs were derived from H9, circle, or HUES4, triangle, ESCs). Data are represented as mean ± SEM (*N*=6). \*\**P*<0.01, \*\*\*\**P*<0.0001 (one-way ANOVA Tukey’s multiple comparison test was applied to analyze differences at day 8, only significant comparisons are shown). **d**, Dot plots showing percentage of G1, S, and G2M cycling cells assayed by flow cytometry with EdU and DAPI staining in control and HHEX knockdown EP expansion. Data are represented as mean ± SEM (*N*=6). \**P*<0.05, \*\**P*<0.01, \*\*\**P*<0.001, \*\*\*\**P*<0.0001 (one-way ANOVA Tukey’s multiple comparison test, only significant comparisons are shown). Representative flow cytometry plots are in Extended Fig. 1e. **e**, Representative images of control (top row) and HHEX shRNA (bottom row) EP cells stained with EdU, FOXA2, HHEX, DAPI. Scale bar = 50 m. **f**, Top row: representative immunostaining of PDX1 and SOX9, including DAPI, of VFG-derived pancreatic spheroids at passage 5. Bottom row: representative immunostaining of AFP and ALB, including DAPI, of VFG-derived hepatic organoids at passage 5. Scale bar = 50µm.

As ventral foregut endoderm in mouse is actively cycling^22^, we measured the proliferation rate of EP culture as a function of time in expansion and found that it increased with time in culture (p6, p8, p12, and p15) (Fig. 1c). In mouse, the TF HHEX is known to support ventral foregut expansion and morphogenesis^21^ and to further confirm the identity of EP we knocked down HHEX by shRNA (Extended Fig. 1d) and observed a reduction in growth. To further quantify this effect, we measured the number of actively proliferating cells in transient endoderm (ADE), endoderm expansion (EP cells) and HHEX knock down EP cells by 5-ethynyl-2’-deoxyuridine (EdU) labelling followed by cell-cycle analysis based on DAPI staining (Extended Fig. 1e). The percentage of cells in S-phase increased with time in expansion, but this was dependent on HHEX, and we observed a reciprocal expansion and HHEX dependent reduction in cells in G2M (Fig. 1, d and e). Based on the expression of ventral foregut markers in EPs, the cytokines used to support these cultures, and function of HHEX in proliferation, we conclude that human EP cells are an *in vitro* model for human ventral foregut and refer to them hereafter as ventral foregut progenitor cells (VFGs).

Ventral foregut gives rise to ventral pancreatic bud, hepatocytes, and the biliary system^23^. Similarly, VFG culture have been shown to produce both pancreatic and hepatic cell types upon differentiation^17^. To further probe VFG differentiation efficiency, we established VFG cultures from a hESC line containing a pancreatic differentiation reporter (PDX1-eGFP)^24^ and determined the minimal cytokine set required to transform VFG spheres into either proliferating pancreatic spheroids or hepatic organoids, focused on the common requirement of BMP antagonism and FGF signalling in these distinct differentiation trajectories (Extended Fig. 1f). Removal of BMP4 from the culture media produced negligible activation of the PDX1 reporter (< 2% eGFP positive, Extended Fig. 1g), no PDX1 protein (Extended Fig. 1h) and no dramatic transcriptional level change at single-cell level (Extended Fig. 1i). The subsequent addition of FGF7 and 10, and to a lesser extent FGF2, produced robust stimulation of PDX1-eGFP (Extended Fig. 1, j and k) and induced robust transcriptional change (Extended Fig. 1i). In response to the initial cytokine treatment, we found we could separate PDX1^+^ (PDX1 positive) and PDX1^-^ (PDX1 negative) cells, and that PDX1^+^ cells expand as pancreatic spheroids, whereas PDX1^-^ cells expand as hepatic organoids (Fig. 1f and Extended Fig. 1, l-n) in defined media^25, 26^.

Taken together these observations indicate human VFG culture is uniquely poised to generate expanding hepatic and pancreatic endoderm.

### Enhanced pancreatic differentiation of VFG cells as a function of time in expansion

To directly compare the differentiation efficiency of expanding VFGs to standard differentiation we employed aspects of three well-established protocols for the differentiation of pancreatic endoderm (PE) derived from hESCs^12, 14, 24^ (Extended Fig. 2a). In two of these protocols^14, 24^ we observed relatively inefficient differentiation, with < 20% of these cultures becoming PDX1-eGFP^+^ (Fig. 2a and Extended Fig. 2b). However, a protocol that couples the inhibition of BMP signalling to stimulation of FGF and WNT signalling^12^, resulted in > 80% PDX1 induction, suggesting that VFG cultures may be best adapted to protocols that harness signals regulating ventral pancreatic specification. VFG-derived PE expressed pancreatic markers including *PDX1* and *NKX6-2*, pancreatic progenitor cell surface marker Glycoprotein 2 (GP2)^24, 27^ and the ventral pancreatic marker Roundabout2 (ROBO2)^28^ (Extended Fig. 1c). Consistent with the observation that ventral pancreatic bud expands more than the dorsal bud^20^, cells differentiated via this third protocol and not the other two, continue to proliferate (Fig. 2, b and c).

**Fig. 2:**
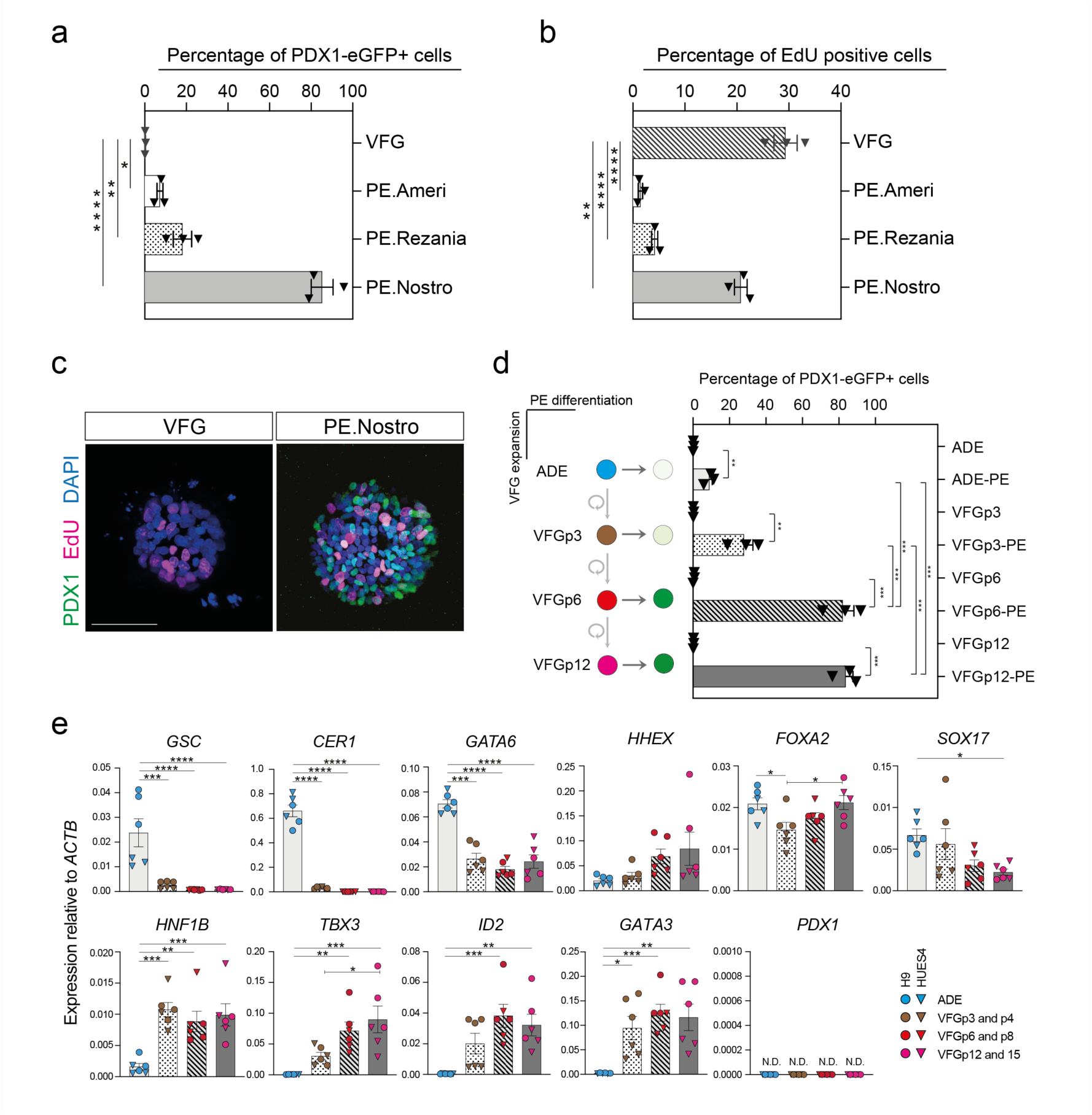
Expansion enhances pancreatic differentiation of VFG cells. **a-b**, Box plots showing percentage of PDX1+ve (a) or EdU+ve (b) cells from flow cytometry analysis in VFG cells, and pancreatic endoderm (PE) generated from VFG cells with protocols from Ameri et al.^24^, Rezania et al.^14^ or Nostro et al.^12^. Data are represented as mean ± SEM (*N*=3). *p<0.05, **p<0.01, \*\*\*\**P*<0.0001 (one-way ANOVA Dunnett’s multiple comparison test compared with VFG cells). **c**, Representative immunostaining of VFG cells and PE, generated using conditions from Nostro et al.^12^, stained with PDX1, EdU, DAPI. Scale bar = 50µm. **d**, Left: schematic representation showing PE differentiation using conditions from Nostro et al.^12^ from ADE, VFG at p3, p6, and p12. Right: box plots showing percentage of PDX1-eGFP positive cells generated for the indicated conditions. Data are represented as mean ± SEM (*N*=3). Statistical analysis was performed between each starting cell type and its differentiation (\*\**P*<0.01, \*\*\**P*<0.001, unpaired two-tailed t-test), as well as between differentiation experiments that start from different cell types (one-way ANOVA Tukey’s multiple comparison test, only significant comparisons are shown). **e**, Expression analysis of VFG culture at different passage numbers: qRT-PCR of primitive streak (*GSC* and *CER1*), pan-endodermal (*GATA6*, *FOXA2*, *HHEX*, *SOX17*), foregut (*HNF1B*) genes, VFG specific (*TBX3, ID2, GATA3*)^17^ and pancreatic progenitors (*PDX1*) across transient ADE and VFG in early (p3 and p4), middle (p6 and p8), and late passage (p12 and p15). Triangles and circles mark cells derived from HUES4 and H9 ESCs respectively. Expression is normalized with *ACTB*. Data are represented as mean ± SEM (*N*=6). *P<0.05, **P<0.01, ***P<0.001, ****P<0.0001 (one-way ANOVA Dunnett’s multiple comparison test compared with ADE, only significant comparisons shown).

To assess the impact of expansion on downstream differentiation, we compared the ability of different passages of VFGs to undergo pancreatic differentiation. Differentiation efficiency increased with time in expansion and was maintained at a similar level following six passages (Fig. 2d and Extended Fig. 2d). To examine why expanding VFG cultures at later passages exhibit enhanced capacity to differentiate, we measured expression of known primitive streak and pan-endoderm/foregut markers, as well as transcriptional regulators reported to be enriched in human endodermal progenitors^17^ during expansion (Fig. 2e). Expression of primitive-streak and early endoderm genes, *GSC, GATA6* and *CER1*, decreased upon expansion. General endoderm markers known to be expressed in the ventral foregut, such as *FOXA2*, *HHEX*, and *SOX17*, were expressed throughout expansion at levels comparable to those in transient ADE cells. Expression of foregut marker, *HNF1B*, together with the ventral foregut markers, *TBX3*, *ID2*, and *GATA3*, were elevated at early passaged VFG cells (p3 and p4) and maintained during expansion. The pancreatic progenitor marker *PDX1* was not detected during VFG expansion at any time point.

### Chromatin accessibility and gene regulation in VFG expansion and onward differentiation

To confirm the stability of the VFG transcriptional state we determined the VFG transcriptome at multiple passages by RNA-seq. Principle component analysis (PCA) showed that VFG cells form a cluster separate from ADE and ESC cells, confirming previous observations^17^ (Fig. 3a). Different passages of VFGs, cultured both with and without BMP4, cluster together and separate in the first principle component from different pancreatic endpoints (Fig. 3a). To explore why expanding VFG cultures exhibit enhanced capacity to differentiate, despite the absence of substantial transcriptional changes during expansion, we determined whether chromatin accessibility changes as a result of prolonged culture. We used ATAC-seq to map chromatin accessibility during the stepwise progression of hESC toward the pancreatic progenitors, at five defined stages of differentiation and expansion: ESC, ADE, VFGp3, VFGp6, and PE. The progression of ESC, through ADE to VFGs and PE is similar to that observed in the RNA-seq (Fig. 3b). However, unlike the transcriptome of different passage VFG cultures that cluster together by PCA, we observed considerable change in the ATAC-seq as a function of time in culture, with the higher passage VFGs moving toward PE (Fig. 3b).

**Fig. 3:**
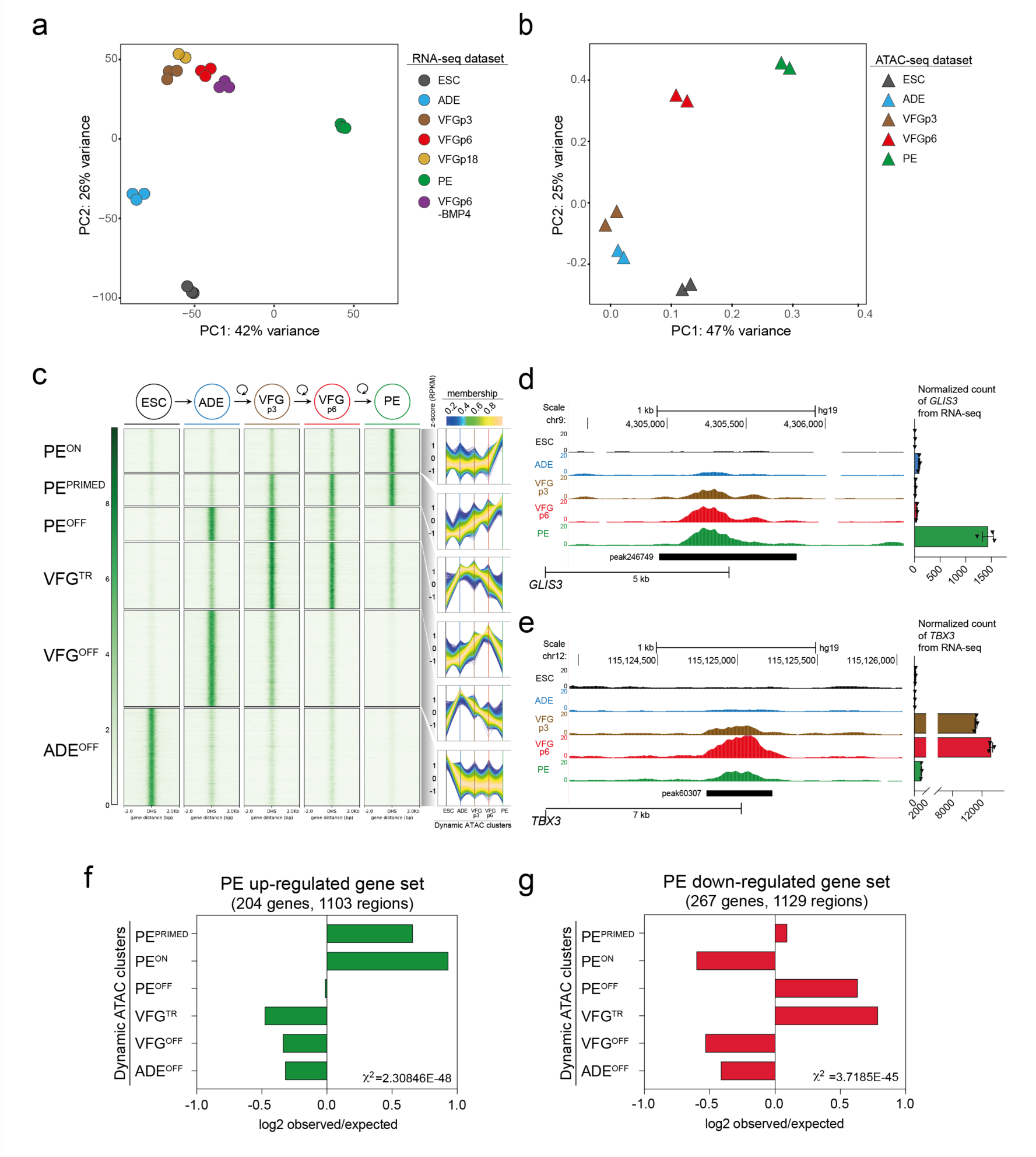
Dynamic chromatin accessibility and gene expression during VFG expansion and pancreatic differentiation. **a**, Principle component analysis (PCA) based on top 2000 differentially expressed genes in bulk RNA-seq dataset of ESC, transient ADE and VFG cells (at p3, 6, and 18), VFG cells without BMP (at p6) and PE cells generated from VFGp6 cells. **b**, PCA of ATAC-seq dataset for ESC, transient ADE and VFG cells (at p3, 6). **c**, Left: Heatmaps of the normalized ATAC-seq signal for the dynamic clusters identified by c-means clustering. Right: Time course-sequencing (TCseq) trajectories for each cluster. Membership score reflects how well a given enhancer follows the pattern identified in time course analysis. The complete set of ATAC-peaks for the dynamic clusters are contained in Supplementary Table 1a. **d-e**, Left: UCSC Genome Browser screen shot at the *GLIS3* (**d**) and *TBX3* (**e**) locus showing ATAC-seq data from ESC, ADE, VFGp3, VFGp6 and PE. Genome coordinates (bp) are from the hg19 assembly of the human genome. The PE^PRIMED^ regulatory element (peak246749) (**d**) and VFG^TR^ element (peak60307) (**e**) are shown with a black bar and the approximate distance between the element and the respective TSS is indicated. Right: RNA-seq data (normalized read count) for *GLIS3* (**d**) and *TBX3* (**e**) across the same conditions as the ATAC tracks. Data are represented as mean ± SEM (*N*=3). **f-g**, Barplot showing the prevalence (log2 observed/expected) of ATAC peaks within a 200Kb window from genes upregulated (**f**) or downregulated (**g**) between PE and VFGp6 across the defined ATAC peak clusters. Genes considered here had a base mean expression > 1000, absolute log2FC > 1.5 and adjusted p-val < 0.05. For annotation see Supplementary Table 1, d and f. Analysis using lower base mean (100) or reduced genomic window sizes (25 Kb) are shown in Extended Fig. 3, e and g. All data shown are significant using chi-square analysis.

In order to identify dynamic changes in open chromatin at regulatory elements that are putative enhancers, we used general linear modelling^29^ to define the dynamic changes in chromatin accessibility at promoter-distal ATAC-seq peaks across these five stages of differentiation. This resulted in a dynamic set of 57803 sites (Extended Fig. 3a) that show significant opening or closing of chromatin in at least one of the stages of differentiation. We defined temporal patterns of chromatin accessibility using c-means clustering which resulted in 8 clusters corresponding to six distinct groups of putative enhancers (Fig. 3c and Supplementary Table 1a). The largest group (ADE^OFF^, 15,291 peaks, 26.5%) correspond to sites where chromatin accessibility is shut down at the start of differentiation from ESCs to ADE and remains closed for the rest of the course of differentiation. The large VFG^OFF^ cluster (15,093 peaks, 26.1%) correspond to sites that become accessible during ESC to ADE differentiation, but that then progressively lose accessibility during VFG differentiation or expansion, so that they are inaccessible in PE. The PE^OFF^ cluster (5234 peaks, 9.1%) also appears at ADE and then loses accessibility but only after VFG expansion. The PE^ON^ cluster (6788 peaks, 11.7%) encompasses regions that only open up during differentiation to PE. We defined two VFG clusters, VFG transient (VFG^TR^) and PE^PRIMED^ clusters.

Chromatin accessibility for the PE^PRIMED^ cluster (n= 4976, 8.6%) increases gradually during VFG expansion and is most accessible in PE. An example of this is an element (peak246749) located ∼5 Kb upstream of *GLIS3* TSS (Fig. 3d, left), and with two regions contained with an ∼200 Kb window spanning the locus (peak246735 and peak246752 in Extended Fig. 3b). Mutations and common genetic variants at *GLIS3* are associated with neonatal diabetes and diabetes risk, respectively. *In vivo Glis3* is expressed in pancreatic endocrine progenitors and then beta cells, where it has an important regulatory function^30^. RNA-seq shows that *GLIS3* is not expressed until PE differentiation from expanded VFGs (Fig. 3d, right). We also observed some increases in accessibility in the conserved enhancer regions (I-IV) of the master pancreatic regulator *PDX1* gene^31, 32^ (Extended Fig. 3c), suggesting that the development of the regulatory landscape at these loci during VFG expansion primes gene activity for later expression in PE. The VFG^TR^ cluster (10,421 peaks, 18% of dynamic peaks) contains regions where chromatin accessibility increases during the VFG expansion, but is then shut down during onward differentiation to PE. Examples of this are found at the *TBX3* putative enhancers located ∼7 Kb upstream of TSS (peak60307) (Fig. 3e, left) and two regions across ∼40 Kb of the *TBX3* loci (peak60300 and peak60310) (Extended Fig. 3d). *TBX3* is expressed in the developing human posterior foregut, and liver bud progenitors ^21, 33^. Consistent with this region driving transient expression of *TBX3* during the differentiation time course, RNA-seq showed that *TBX3* expression is induced during the differentiation of ADE to VFGp3, is maintained during VFG expansion, but then lost during the differentiation to PE (Fig. 3e, right).

To link these clusters of dynamic enhancers with changes in gene expression we defined a list of genes that significantly change their expression in the transition from expansion into further differentiation (log2 fold change > 1.5, *p*-val. < 0.05) (Supplementary Table 1b). To pair enhancers with specific genes, we considered dynamic enhancers located either within 25 Kb or 200 Kb of the single nearest gene’s transcriptional start site (TSS) (Supplementary Table 1a) and we excluded low level changes in basal gene expression (Supplementary Table 1, c and d). While filtering out gene expression noise that occurs with passaging reduces the size of the gene set, we defined 1103 dynamic enhancers located within 200kb of 204 up-regulated PE genes (Supplementary Table 1d) (compared to 367 dynamic enhancers for 65 genes, when a 25 Kb cut off is used). Regardless of which enhancer set we used, we observed significant enrichment of both PE^PRIMED^ and PE^ON^ enhancer classes with up-regulated PE genes (Fig. 3f and Extended Fig. 3, e and f, although the enrichment is greater for enhancers located in closer proximity to the genes they regulate. We also identified regulatory regions within 200kb of genes down regulated in differentiation with 267 down-regulated genes associated with 1129 dynamic enhancers (Supplementary Table 1, e and f) (462 dynamic enhancers for 83 genes at 25 Kb). These down-regulated PE gene sets were associated with the PE^OFF^ and VFG^TR^ enhancer categories (Fig. 3g; Extended Fig. 3, g and h). Taken together, this suggests that VFG expansion primes some pancreatic enhancers for later induction of their target genes in differentiation and decommissions enhancers inappropriate for the pancreatic endoderm lineage, insuring that their targets will not be inadvertently expressed in this lineage.

### VFG expansion enables consolidation of an enhancer landscape that is imperfectly realized during directed differentiation

We next wished to understand the extent to which the enhancer network induced during expansion is normally exploited in directed differentiation. A previous study used ATAC-seq to profile chromatin accessibility changes during the differentiation of hESCs through definitive endoderm (DE), and posterior foregut (FG) stages to pancreatic progenitors (PP1)^34^. Comparing the behaviour of ATAC-seq peaks from those data across the six differentiation patterns we have classified from our culture system, we can define a common set of putative enhancers activated either in VFGs (this study) or FG^34^ and that then remain accessible in the final differentiation stage (PE or PP1), respectively (PE-PP1 common, Fig. 4a and Extended Fig. 4a; Supplementary Table 2a); and a class of element that fails to be induced in the absence of expansion (PE-not-PP1, Fig. 4a and Extended Fig. 4a; Supplementary Table 2a). An example of this is an element located ∼66 Kb downstream of the *FGFR2* TSS (peak35254) (Fig. 4b). Many of the peaks defined as pancreatic endoderm elements in Lee et al.^34^, correspond with peaks that we find are closed down during or after VFG expansion (VFG^OFF^-in-DE-PP1 and VFG^TR^-in-PP1, Fig. 4a and Extended Fig. 4a; Supplementary Table 2a). An example of this is an element located ∼68 Kb downstream of the *MEF2C* TSS (peak192828) (Fig. 4c). Motif analysis reveals an enrichment of TFs relevant to pancreatic and liver development in human embryos^35, 36^, such as ONECUT, FOXA, TEAD, and HNF1B, in both the PE-not-PP1 and the PE-PP1 common clusters (Extended Fig. 4b). Enriched TF motifs for VFG^TR^-in-PP1 and VFG^OFF^-in-DE-PP1 do not have a strong link to pancreatic-specific differentiation or function. Together, these data suggest that VFG expansion allows for the commissioning of enhancers relevant to pancreatic differentiation and the decommissioning of enhancers for alternative lineages. This process appears to be bypassed in directed differentiation.

**Fig. 4:**
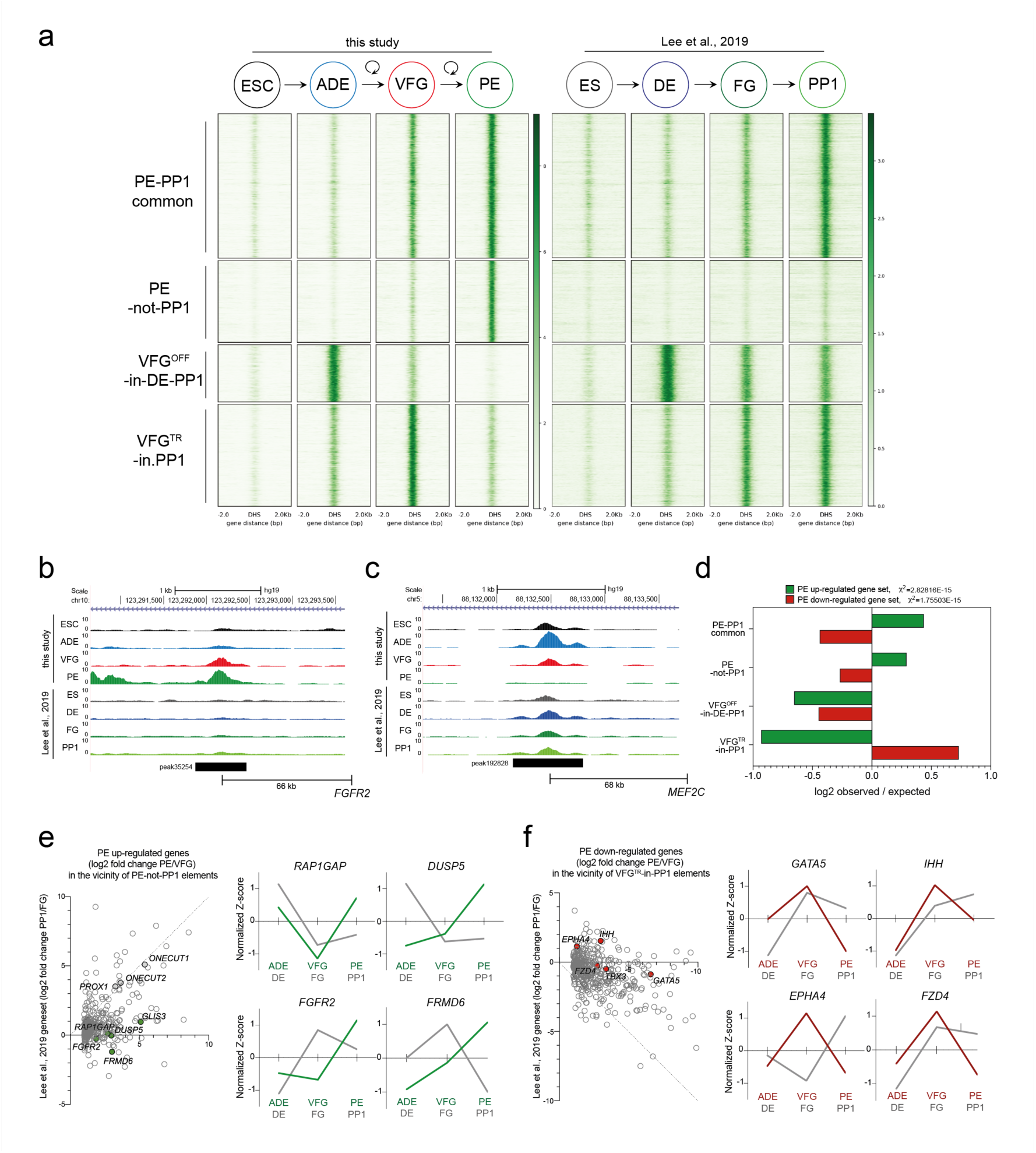
VFG expansion enables consolidation of an enhancer landscape that is imperfectly realized during directed differentiation. **a**, A comparison of chromatin accessibility of enhancers charted in this study (Heatmap, left) with the Lee et al. dataset^34^ (Heatmap, right). Enhancers in the group “PE-PP1 common” are the pancreatic endoderm enhancers that are activated independent of VFG expansion (n=7504). Enhancers in the group PE-not-PP1 are PE enhancers that are activated only if PE is differentiated from expanding VFGs (n=4260). Enhancers in the “VFG^OFF^-in-DE-PP1” group, represent a subset of ADE enhancers that are inactivated during VFG expansion (n=2974). The “VFG^TR^-in-PP1” enhancer group at the bottom of the heatmap (n=5293) are inactivated in PE derived from expanding VFGs, but not in directed differentiation. **b-c**, UCSC browser screenshot showing examples of a PE-not-PP1 enhancer (peak35254), in an intron of the *FGFR2* locus (**b**) and a VFG^OFF^-in-DE-PP1 group (peak192828) contained with in an intron of the *MEF2C* locus (**c**). Approximate distance between elements and TSS is indicated. A complete set of VFG-specific ATAC-clusters is contained in Supplementary Table 2a. **d**, Barplot showing the prevalence (log2 observed/expected) of ATAC peaks within a 200Kb window from genes up-regulated (green) or down-regulated (red) between PE and VFGp6 across the defined ATAC peak clusters. Genes considered have a base mean expression > 1000, absolute log2FC > 1.5 and adjusted p-val < 0.05. For annotation see Supplementary Table 2, b and c. All data shown are significant in chi-square analysis. **e-f**, Left: Scatter plot of gene expression for genes up-regulated (**e**) or down-regulated (**f**) in PE vs VFG (as defined in d), and within 200Kb of minimum one ATAC peak in the PE-not-PP1 (**e**) or VFGTR-in-PP1 (**f**) clusters respectively, vs their expression after directed differentiation (PP1/FG). The diagonal line indicates where there is no difference in differential expression between two comparisons (datasets). Right: normalized z-score expression of representative candidates (**e**: RAP1GAP, DUSP5, FGFR2, FRMD6 and **f**: GATA5, IHH, EPHA4, FZD4). Normalized z-score expression for each candidate was plotted for the ADE, VFG (p6), and PE conditions (green), and the DE, FG, and PP1 conditions (grey).

Mapping of these enhancer elements to potential target loci (based on positions with 200 Kb of the nearest gene) (Supplementary Table 2, b and c), reveals an enrichment for the two pancreatic endoderm enhancer clusters, PE-PP1 common and PE-not-PP1, in the vicinity of genes up regulated in VFG-derived pancreatic endoderm (the same gene set used for Fig. 3, f and g) (Fig. 4d). However, when we explored elements induced in directed pancreatic differentiation, but not active in VFG-derived PE, our PE up-regulated gene set did not correlate with either the VFG^OFF^-in-DE-PP1 or VFG^TR^-in-PP1 classes of elements, although the PE down-regulated gene set correlates with location of VFG^TR^-in-PP1 elements (Fig. 4d), suggesting expansion decommissions enhancers that could be driving lineage non-specific gene expression.

To explore global correlations between genes differentially regulated in pancreatic endoderm derived from expanding VFGs and directed differentiation, we plotted gene expression in both protocols for the two classes of expansion dependent elements, PE-not-PP1 (Fig. 4e, left) and VFG^TR^-in-PP1 (Fig. 4f, left; Supplementary Table 1, d and f). Expression of genes in the vicinity of PE-not-PP1 elements are better induced in VFG-derived PE than directed differentiation, whereas, genes mapped to elements decommissioned in expansion, VFG^TR^-in-PP1, are more extensively down regulated when PE is differentiated from expanding VFGs. Expansion dependent positive and negative gene regulation can also be seen when the expression of individual mRNAs is compared, where we observed expansion dependent up regulation of *RAP1GAP*, *DUSP5*, *FGFR2* and *FRMD6* (Fig. 4e, right); and ectopic expression of *GATA5*, *IHH*, *EPHA4*, and *FZD4* (Fig. 4f, right) in the directed differentiation. Taken together, this analysis suggests that there are significant differences in mRNA expression related to expansion dependent changes in enhancer accessibility.

### VFG expansion captures enhancers active during organogenesis of the human ventral foregut

To determine how the dynamic enhancer landscape we have captured during VFG expansion and PE differentiation *in vitro* correspond with the regulatory landscape during *in vivo* pancreatic development, we compared our ATAC-seq data with H3K27ac data obtained from micro-dissected endodermal (pancreatic, liver, and lung), mesodermal (adrenal and heart), and ectodermal (RPE and brain) tissues collected from Carnegie stages 15-22 human embryos^37^ (Supplementary Table 3). Consistent with the VFG identity of these expanding cultures, the PE^PRIMED^ class of element is enriched in both liver and pancreatic enhancers, while the PE^ON^ class is more enriched in pancreatic elements (Fig. 5a and Supplementary Table 4a). Examples of these are found spanning a combination of PE^PRIMED^ (peak 97567) and PE^ON^ (peak 97566) elements located in the first intron of *HNF1B* gene (Fig. 5b). In mouse, Hnf1b is a key member of the network of transcription factors controlling the differentiation of pancreatic multipotent progenitor cells to acinar, ductal and endocrine^38^, that is also necessary for the induction of liver organogenesis and for the activity of mature hepatocytes^39, 40^. Enhancer clusters that shut down as expanded VFGs differentiate to PE (VFG^TR^ and PE^OFF^) are most enriched for enhancers active in the developing liver, consistent with differentiation away from the pancreatic lineage (Fig. 5a and Supplementary Table 4a). Elements that lose their accessibility early in our differentiation or expansion (ADE^OFF^ or VFG^OFF^) are most enriched for predicted enhancers active in the ectodermal and mesodermal tissues consistent with the endodermal identity of VFG culture (Fig. 5a and Extended Fig. 5a; Supplementary Table 4, a and b).

**Fig. 5:**
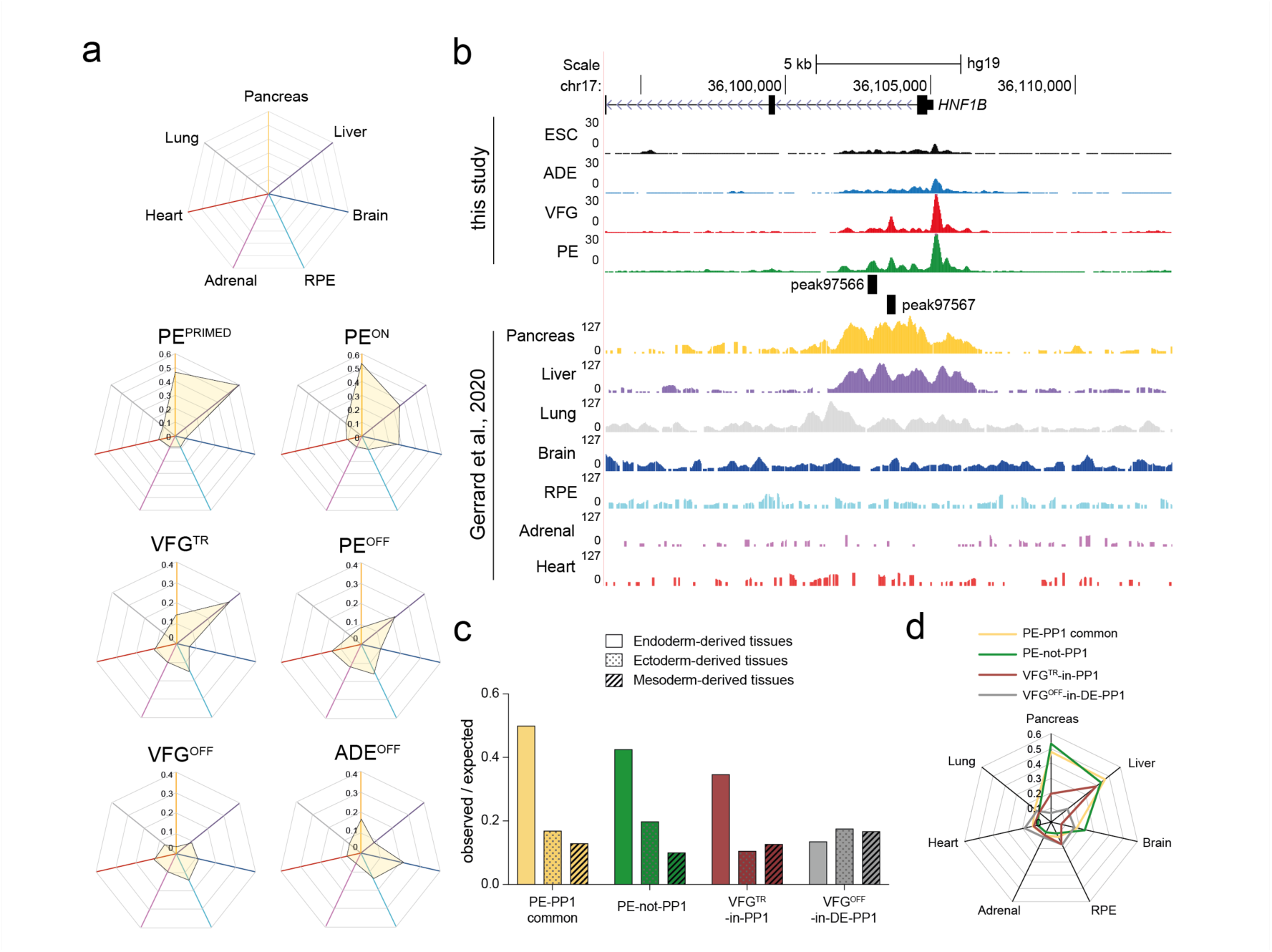
VFG expansion captures enhancers that are active during human ventral foregut derived organogenesis. **a**, Enrichment of tissue-specific H3K27ac enhancers from human embryos in different ATAC clusters defined in Fig. 3c were displayed by enrichment score (observed/expected) in radar charts. A complete set of lineage and tissue specific H3K27ac enhancer is contained in Supplementary Table 3 and their overlaps with ATAC-clusters in Supplementary Table 4, a and b. **b**, UCSC Genome Browser screen shot at the *HNF1B* locus showing ATAC-seq data from this study (ESC, ADE, VFG, and PE) and H3K27ac ChIPseq data (Hanley Lab 2014, 2015, STAR_hg38_FULL, ds99 scaled)^37, 78^ from multiple human embryonic tissues (pancreas, liver, lung, brain, RPE, adrenal, and heart). Genome coordinates (bp) are from the hg19 assembly of the human genome. PE^PRIMED^ (peak97567) and PE^ON^ (peak97566) elements overlapping with pancreatic specific H3K27ac enhancer are shown at the bottom and the approximate distance between the elements and the *HNF1B* TSS is indicated. **c**, Enrichment of lineage-specific H3K27ac enhancers (endoderm, ectoderm, and mesoderm) from human embryos^37^ in the different VFG-specific ATAC clusters defined in fig. 4a by enrichment score (observed/expected). **d**, Enrichment of tissue-specific H3K27ac enhancers from human embryos across different VFG-specific ATAC clusters defined in fig. 4a by enrichment score (observed/expected) in a radar chart. A complete set of lineage and tissue specific H3K27ac enhancer overlapped with VFG-specific ATAC-clusters is listed in Supplementary Table 4c.

As VFG expansion and differentiation directly from pluripotent cells are both *in vitro* models for aspects of human development, we sought to define the extent to which the difference we observed in sets of dynamic enhancer classes compare to those associated with human organogenesis. We found that the expansion dependent PE-not-PP1 class of element was enriched in enhancers common to early fetal endoderm and more specifically the ventral foregut derived pancreas and liver (Fig. 5, c and d; and Supplementary Table 4c). An example of this is an element (peak60959 in Extended Fig. 5b) located in the first intron of *HNF1A*, one of the most common causes of maturity-onset diabetes of the young (MODY) diabetes^41^. Moreover, the set of elements that lose their chromatin accessibility during expansion (VFG^OFF^-in-DE-PP1) or in VFG differentiation to PE (VFG^TR^-in-PP1), did not contain significant numbers of pancreatic elements, but represent a combination of liver (with an example shown at *HNF4A* locus, peak142009 in Extended Fig. 5c) and other organs (Fig. 5, c and d; and Supplementary Table 4c).

### FOXA proteins are required for VFG enhancer priming in pancreatic differentiation

To determine factors responsible for VFG enhancer priming we assessed the enrichment of specific TF motifs in different dynamic enhancer classes (Fig. 6a and Supplementary Table 5). The PE^ON^ enhancer class are strongly enriched in ONECUT1 and two factors previously associated with pancreatic differentiation CUX1 and CUX2^42^. In contrast, the PE^PRIMED^ enhancer class are enriched in FOX motifs (in particular FOXA) and to a lesser extent a number of unrelated endodermal/hepatic factors broadly classed as hepatic nuclear factors (HNFs)^43^ (Fig. 6a). Both enhancer classes were enriched in binding sites for CTCF and TEAD factors. To further refine the association of specific TFs with these enhancer classes, we used k-means clustering to define patterns of mRNAs expression associated with enhancers that are up regulated in pancreatic endoderm (Extended Fig. 6a). Focusing on patterns of gene expression that increase in PE relative to VFG culture for genes associated with PE^PRIMED^ enhancers (K-clusters 3, 8, and 10; Extended Fig. 6b, left), we identified 1150 regions enriched in FOX motifs, and a similar spectrum of TFs to those shown in Fig. 6a. Similar analysis of the PE^ON^ (K-clusters 1, 4, and 6) enhancers shows enrichment in CUX and FOX motifs but not HNF1B (Extended Fig. 6b, right). Of all the FOX factors known to recognize these DNA binding sites, a combination of embryonic expression and phenotypes in mouse development suggest FOXA1 and 2 could be relevant to VFG mediated enhancer priming^32, 44^. FOXA2 is required for both pancreas development in mouse^32^ and human ESC differentiation^34^, and the role of FOXA1 in pancreas development becomes apparent in the context of compound FOXA1/2 double mutants. FOXA factors are known ‘pioneer factors’ that access regulatory regions and prepare them for later activation^45^, but FOXA1 mutant ESCs exhibits no defect in directed pancreatic endoderm differentiation^34^.

**Fig. 6:**
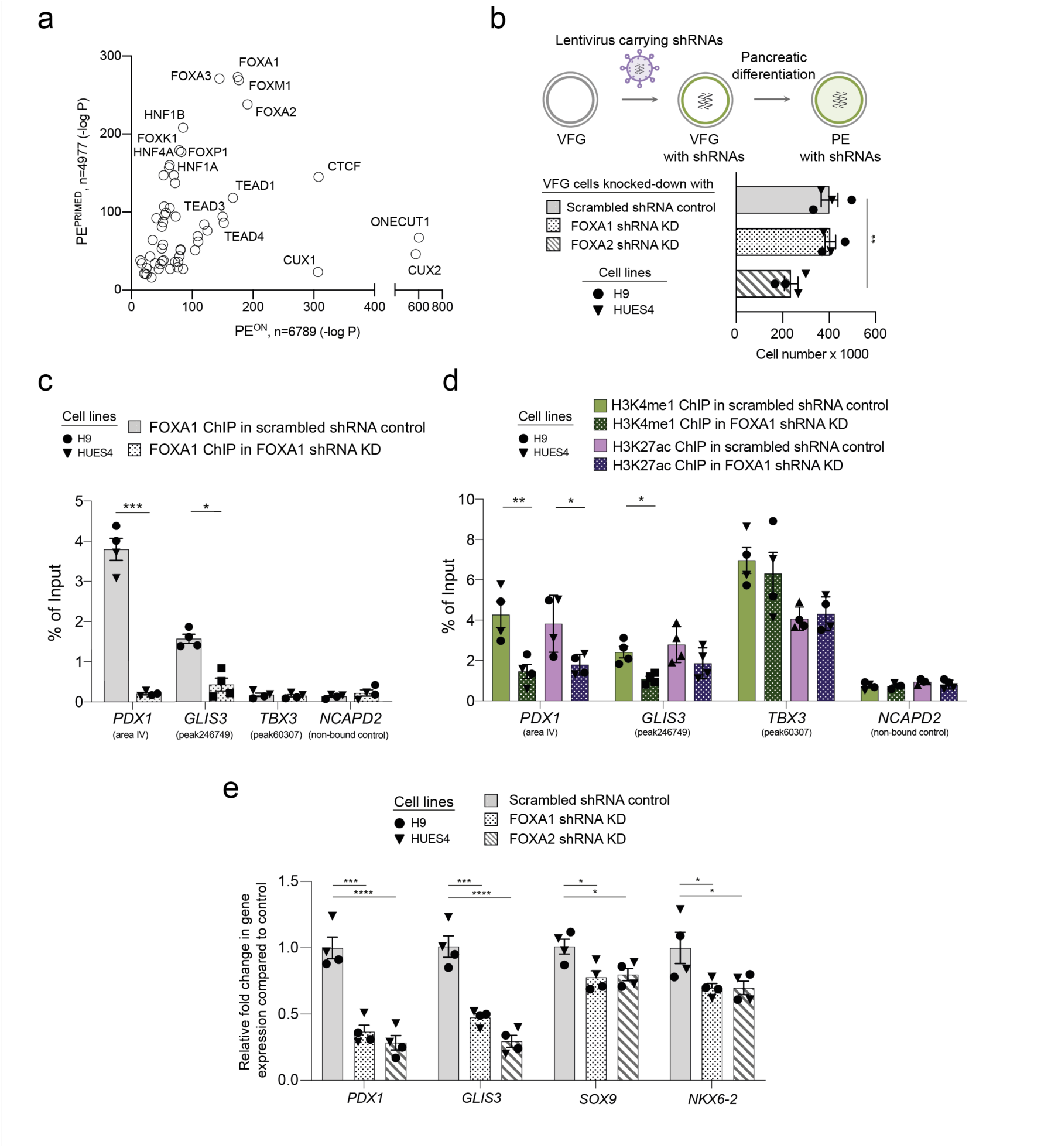
FOXA proteins are required for VFG enhancer priming towards pancreatic differentiation. **a**, Transcription factor motif enrichment in PE^PRIMED^ and PE^ON^ ATAC clusters. n = peaks analysed. *P* values were derived from hypergeometric enrichment using HOMER default background (see Supplementary Table 5, a and b). Candidate factors with *P*-value > 1e^-10^ were not included in the plot. **b**, Knock-Down (KD) of FOXA1 and FOXA2 in both H9- and HUES4-derived VFG cells. Top: Schematic of generation of FOXA1 and FOXA2 shRNA KD VFG cells and PE differentiation of these. Bottom: Box plot for proliferation assay (cell counts) for FOXA1 and FOXA2 shRNA KD and scrambled shRNA control VFG cells. Data are represented as mean ± SEM (*N*=4). Statistics analysis was performed between KDs and control VFG cells (\*\**P*<0.01, unpaired two-tailed t-test, only significant comparisons are shown). **c**, FOXA1 binding enrichment by ChIP-qPCR at enhancer regions of *PDX1* (area IV), *GLIS3* (peak246749), and *TBX3* (peak60307) in FOXA1 shRNA KD VFG and scrambled control cell lines. An intragenic region of *NCAPD2* served as non-bound control. Data are represented as mean ± SEM (*N*=4). Statistics analysis was performed between KD and control VFG cells (\**P*<0.05, \*\*\**P*<0.001, unpaired two-tailed t-test, only significant comparisons are shown). **d**, H3K4me1 and H3K27ac enrichment by ChIP-qPCR at enhancer regions of *PDX1* (area IV), *GLIS3* (peak246749), and *TBX3* (peak60307) in FOXA1 shRNA KD VFG and scrambled control cell lines. An intragenic region of *NCAPD2* served as a non-bound control. Data are represented as mean ± SEM (*N*=4). Statistics analysis was performed between the KD and control VFG cells (\**P*<0.05, \*\**P*<0.01, unpaired two-tailed t-test, only significant comparisons are shown). **e**, Differentiation of FOXA1 and FOXA2 shRNA KD and scrambled control VFG cells to PE. Relative fold change in mRNA of pancreatic genes (*PDX1*, *GLIS3*, *SOX9* and *NKX6-2*) was assayed by qRT-PCR. Expression is normalized to *ACTB*. Data are represented as mean ± SEM (*N*=4). \**P*<0.05, \*\*\**P*<0.001, \*\*\*\**P*<0.0001 (one-way ANOVA Dunnett’s multiple comparison test compared with control).

To test whether FOXA1 and FOXA2 are important for pancreatic priming during human VFG expansion we knocked them down by shRNA during VFG expansion (Fig. 6b and Extended Fig. 6c). Neither factor produced a significant reduction in the expression of VFG markers (Extended Fig. 6, d and e), although FOXA2, but not FOXA1, KD impaired VFG expansion (Fig. 6b). By ChIP-qPCR, we confirmed the binding of FOXA1 in the PE^PRIMED^ enhancers of *PDX1* and *GLIS3* (regions are indicated in Extended Fig. 3c and Fig. 3d), but not in the VFG^TR^ element of *TBX3* and that this binding was reduced in the stable FOXA1 KD cell lines (Fig. 6c). ChIP-qPCR for histone modifications associated with enhancer priming, H3K4 mono-methylation (H3K4me1) and H3K27 acetylation (H3K27Ac) shows that knock down of FOXA1 leads to a significant reduction in these chromatin marks in PE^PRIMED^ enhancers associated with *PDX1* and *GLIS3* (Fig. 6d), but not VFG^TR^ elements in *TBX3*. When either FOXA1 or FOXA2 KD VFG cells were challenged in pancreatic differentiation, expression of the pancreatic markers was significantly reduced (Fig. 6e). Taken together this analysis suggests that VFG expansion enables pioneer FOXA factors to prepare enhancer regions for later activity. The role of FOXA1 is particular interesting, as it is not required in the directed differentiation of human ESCs to pancreatic endoderm^34^.

## Discussion

In this paper we show that proliferation of lineage-restricted progenitors enables cells to prepare for onward differentiation. We found that expansion of pluripotent stem cell derived endodermal progenitors that acquire VFG identity primes their enhancer network for organ specific differentiation to the ventral foregut associated lineages of pancreas and liver. Consistent with this, we found that endodermal pioneer factors such as FOXA1 and 2 are required for expansion dependent priming of downstream differentiation, consistent with their role in development of the liver and pancreas^46^.

A portion of the pancreas comprising the uncinate process, in addition to the liver and gall bladder, are all derived from the VFG region of the developing embryo beginning at E8.5 in mouse or at Carnegie stage 10 (25-27 days post coitum) in human^20, 23^. Based on gene expression and differentiation competence, we have identified expanding hESC-derived EP cells as VFGs, suggesting they represent a human cell culture model for this region of the embryonic gut. While prior studies have shown that VFG expansion can produce functional pancreatic endocrine cells^17^, here we demonstrate that this is a direct consequence of time in VFG culture and the resulting equilibration of the enhancer network. *In vivo* pancreas development begins from two independent locations, the dorsal and ventral foregut, induced by different context dependent signals. The dorsal pancreatic endoderm is induced by factors derived from the notochord and dorsal aorta (retinoic acid, activin, and FGF2)^47^, while the ventral pancreatic endoderm is induced in the VFG in the absence of signals driving hepatic specification (FGF2 produced by the cardiac mesoderm and BMP4 originating in the septum transversum)^46^. Embryonic explants from this region appear to contain multi-potent progenitors, but in the absence of exogenous signalling default to pancreas differentiation *ex vivo*^7^. However, *in vivo*, VFG progenitors and their descendants retain multi-potency up to E11.5 in mouse at the base of the three VFG organs^9^. As the location of these progenitor cells is close to both the cardiac mesoderm (FGF source) and septum transversum mesenchyme (BMP source), both components in VFG culture media, it is possible that these founder populations persist via self-renewing cell division in the embryo and that their proliferation may be required for efficient onward differentiation. How many rounds of proliferation are required in these lineage-restricted progenitors to insure high fidelity differentiation is hard to approximate, although the cell cycle length in the foregut endoderm has been estimated based on BrdU labelling as 17.3-26.6 hours depending on the precise location of the progenitor cells^9^. Proliferation in this region *in vivo* is dependent on the homeobox HHEX^22^, similar to the phenotype we observe in response to knocking this factor down in VFG.

TFs are traditionally associated with the recruitment of RNA Polymerase and other components to enhancers and promoters to drive transcription. However, an increasing set of TFs also has the ability to bind DNA in chromatin and destabilise nucleosomes. These pioneer factors include the FOXA proteins identified here as important for VFG priming. In particular, FOXA proteins have been associated with enhancer priming during foregut development^46^ and have the capacity to associate with their enhancers in mitotic chromatin^48^. Yet, FOXA1 is not required for directed differentiation to pancreatic endoderm *in vitro*. One implication of this finding is pioneer activity and, in this instance, FOXA1, depends on proliferation, suggesting that expression of these factors across multiple cells cycles enables the network to physically equilibrate and prepare enhancers for roles in later transcriptional regulation. In the case of FOXA1 in human ESCs, the directed differentiation protocol that comes closest to reproducing the proliferative nature of early ventral foregut, is the one instance where a role for FOXA1 was previously suggested^49^.

Although we are not aware of many examples of a direct comparison of directed differentiation to expansion, it is clear that distinct stages of development exhibit context dependent signalling responses to the same pathways. For example, distinct stages of pluripotency respond to the same cytokines to produce distinct stage specific endoderm^50^, and while the extent to which the reorganization of the enhancer network from naïve to primed pluripotency^51^ requires proliferation has not been explored, the cell cycle at this stage can be as short as five hours^52, 53^ suggesting that cells exhibiting this change in context dependent signalling at these stages could be exploiting cell division even in directed differentiation. Moreover, in both naïve and primed pluripotency the binding of pluripotency TFs to differentiation specific genes can determine how these enhancers will respond to signalling and whether states of differentiation retain plasticity^54, 55^ suggesting that TFs, as we describe here for the FOXA proteins, may have a general role in determining the enhancer sets that will respond to appropriately to lineage specific cues and determine the range of cell types available to different progenitors. In addition to preparing enhancers for later activation, we also found that enhancer decommissioning depends on expansion.

Perhaps these elements are being shut down over time as a result of going through multiple rounds of replication in the absence of specific TFs that can protect these enhancers from nucleosome occlusion following replication. In VFGs, these decommissioned elements were enriched for motifs of the general endoderm inducers GATA4, 5 and 6, suggesting that expansion also shields VFGs from the lingering potential action of early endoderm enhancers during later development. While FOXA1 can bind mitotic chromatin, GATA factors are only partially retained^56^, suggesting that expansion could provide FOXA proteins with a competitive advantage.

At the level of both enhancer commissioning and decommissioning, we observe that the proliferation or expansion as a lineage-restricted progenitor may be essential for high efficiencies in later differentiation. Thus proliferation in embryonic development is not just about producing sufficient numbers of cells, but fine-tuning the response of these cells to upcoming differentiation cues. As it has been suggested that progenitor cell expansion can improve the differentiation efficiency of specific human pluripotent cell lines that perform poorly in specific protocols^18, 57, 58^, perhaps the lineage potential of different pluripotent cell lines is determined by the propensity to proliferate in culture. Moreover, as proliferation and growth are a hallmark of later fetal development, perhaps additional expansion steps could enhance the efficiency with which more mature organ specific cell types can be obtained from human pluripotent cells.

## Materials and Methods

### Experimental Design

#### Maintenance of hESC

Undifferentiated hESCs H9 (WA09, WiCell Madison, WI) were maintained on tissue culture plates pre-coated with 0.1% gelatine with irradiated C57BL6 mouse embryonic fibroblast (MEF) feeder cells (25,000 cells/cm^2^) in H9 hESC media: DMEM/F12 GlutaMAX medium (Thermo Fisher) supplemented KnockOut Serum Replacement (KSR) (Thermo Fisher Scientific), NEAA, beta-mercaptoethanol (Sigma), and 10 ng/mL FGF2 (Peprotech). Cells were passaged as clusters with collagenase IV (Sigma) when reaching approximately 70% confluence, and maintained in 20% O_2_/5% CO_2_/37°C. Undifferentiated hESC HUES4 WT and PDXeG clone 170-3^24^ were adapted and maintained in DEF-CS (Takara). When reaching approximately 80% confluence, cells were dissociated with TrypLE (Thermo Fisher), counted with the automated cell counter NucleoCounter NC-200 (Chemometec). Cells were re-plated at a density of 40,000 cells/cm^2^ and maintained in 20% O_2_/5% CO_2_/37°C. All hESC lines were routinely screened for mycoplasma, and all were negative. All cell lines were approved for use in this project by De Videnskabsetiske Komiteer, Region Hovedstadenunder number H-4-2013-057.

#### Transient differentiation of ADE cells

Transient ADE cells were generated from wild type H9 and HUES4 WT hESCs, as well as HUES4 PDX1-eGFP reporter (PDXeG clone 170-3) hESC cell line^24^ using as described in Cheng et al.^17^. In brief, the hESC cells at 70-80% confluence were collected with Accutase (00455556, Thermo Fisher Scientific), re-plated at a density of 50,000 cells/cm^2^ on a polystyrene cell culture plates (Corning, 353047) pre-coated with un-diluted growth factor reduced Matrigel (GRF-Matrigel; Corning), cultured in either H9 hESC or DEF-CS media for 48 hours with 10µM ROCK inhibitor (Y-27632, STEMCELL Technologies) for first 24 hours and maintained in 20% O_2_/5% CO_2_/37°C. The hESC clusters were used to generate transient ADE cells in three-dimensional differentiation under hypoxic conditions (5% O_2_/5% CO_2_/37°C) for 5 days. In day 1, the cell clusters were cultured in RPMI 1640 GlutaMAX (61870044, Gibco) with 10% SFD media supplemented with Activin A [100 ng/mL], CHIR99021 [3 µM] and 4.5x10-4 M Monothioglycerol (Sigma). On day 2, the medium was changed to RPMI 1640 GlutaMAX supplemented with Activin A [100 ng/mL], BMP4 [0.5 ng/mL], FGF2 [10 ng/mL], VEGF [10 ng/mL], 0.5 mM ascorbic acid [Wako], and 4.5x10^-4^ M Monothioglycerol (Sigma). The same media was applied at day 3. At day 4, differentiation media was changed to SFD media supplemented with Activin A [100 ng/mL], BMP4 [0.5 ng/mL], FGF2 [10 ng/mL], VEGF [10 ng/mL], 0.5 mM ascorbic acid (Wako), and 4.5x10^-4^ M Monothioglycerol (Sigma).

#### Generation and expansion of VFG

EP/VFG expansion was performed as described^17^ with minor modifications. In brief, day 5 transient ADE clusters were dissociated 1 volume of trypsin-EDTA [0.25%] (Thermo Fisher) for 5 minutes at 37°C, and then the enzyme was inactivated with 0.5 volume of fetal bovine serum (FBS; Sigma). Single cells suspension was obtained by repeatedly washing with 10 volumes of ice-cold washing buffer (PBS-/- 3% FBS). The single cells were incubated with 1:100 CXCR4-PEcy7 and CD117-APC (BD Biosciences) for 45 minutes at 4°C and cells were stained with DAPI to exclude dead cells. The CXCR4-CD117 double positive cells were obtained by Fluorescence-Activated Cell Sorting (FACS) on a SH800 (SONY) into SFD media with 1:100 Penicillin-Streptomycin. The sorted cells were re-plated at a density of 20,000-30,000 cells/cm^2^ on polystyrene cell culture plates pre-coated with un-diluted growth factor reduced Matrigel (GRF-Matrigel; Corning) and pre-seeded with low density (8,000 cells/cm^2^) irradiated DR4 mouse embryonic fibroblast (MEF) feeder cells (Lonza) (Matrigel-MEF). Cells were cultured in complete EP/VFG media (SFD medium supplemented with BMP4 [50 ng/ml], FGF2 [10 ng/ml], VEGF [10 ng/ml], EGF [10 ng/ml], 0.5 mM ascorbic acid (Wako), and 4.5x10^-4^ M Monothioglycerol (Sigma)) and maintained under hypoxic conditions (5% O_2_/5% CO_2_/37°C).

The medium was changed every other day until cells reached confluence, at 80,000-120,000 cells/cm^2^. When VFG cells reached approximately 100µm in diameter, they were passaged by dissociation using 1 volume of the trypsin-EDTA dissociation buffer for 5 minutes at 37°C, detached from the plate using a cell scraper and then supplemented with 0.5 volume of FBS for enzyme inactivation. Single cells suspension was obtained by repeatedly washing with 10 volume of ice-cold washing buffer. VFG single cells were re-plated on the pre-coated matrigel with feeders at 15,000-20,000 cells/cm^2^.

#### Single cell preparation for RNA-seq and index sorting

Dissociated ADE and VFG single cells with treatments (mock, BMP4 withdrawal, and BMP4 withdrawal plus FGF2 stimulation) were incubated with 1:100 CXCR4-PEcy7 and CD117-APC (BD Biosciences) for 45 minutes at 4°C and cells were stained with DAPI to exclude dead cells. The single cells from BMP4 withdrawal plus FGF2 stimulated VFG culture were incubated only with 1:100 CD117-APC (BD Biosciences) in a similar condition to that described above. Cells were sorted using a BD FACS Aria III with a 100µm nozzle and 20psi sheath pressure. A forward scatter (FSC) and side scatter (SSC) were used to define a homogeneous population. FSC-H/FSC-W gates were used to exclude doublets and dead cells were excluded based on DAPI inclusion. The boundary between positive and negative populations were set based on negative population of unstained cells. Sorting speed was kept at 100-300 events/s to eliminate sorting two or more cells into one well. Single cell sorting was verified colorimetrically based on a protocol described in Rodrigues and Monard, 2016^59^. Cells were sorted directly into lysis buffer containing the first RT primer and RNase inhibitor, immediately frozen and later processed by the MARS-seq1 protocol as described previously^60^. All single-cell RNA-seq libraries were sequenced using Illumina NextSeq 500 at a median sequencing depth of 225 000 reads per single cell.

#### Immuno-histochemical analysis

Media was removed completely and matrigel dome containing 3D clusters were gently mixed with fresh undiluted matrigel 1:1 and transferred to 8-well µ-slides (Ibidi) wells (20ul/cm2 well) for whole mount immunostaining. When the matrigel was solidified at 37°C, 4% paraformaldehyde (PFA) (at room temperature) was added and cultures were fixed at room temperature for 10 mins, blocked and permeabilized with 2% donkey serum, 0.3% Triton X-100 and 0.1% BSA in PBS for 1 hour at RT. Primary antibodies (Supplementary Table 6) were incubated with 3% FBS in PBS overnight at 4°C and secondary antibodies were added for 1 hour at RT. Brightfield imaging of cells was performed using a Nikon microscope. Fluorescent imaging was done using either a Leica AF6000 fluorescent microscope or a Leica SP8 confocal microscope.

#### EdU labelling

Cells were incubated with 10 µM EdU (Click-iT EdU kit; Thermo Fisher) with medium for 4 hours at 5% O_2_/5% CO_2_/37°C. The 3D clusters were prepared for whole mount immunostaining as described above. Dissociated cells were collected for flow cytometry as described above. Permeabilisation, blocking and Click-iT reaction for EdU detection were performed according to the manufacturer’s instructions. Immunostaining of EdU-labeled 3D clusters were performed with antibodies supplied with the kit and with DAPI (1 µg/ml) for nuclear staining. Flow cytometry of EdU-labeled dissociated cells was performed with DAPI (10 µg/ml) staining cells for DNA content.rede4

#### Flow cytometry

For surface marker staining, dissociated cells were incubated with conjugated antibodies for 1 hour at 4°C and were stained with DAPI (1 µg/ml) to exclude dead cells. For intracellular staining, cells were stained with Ghost Dye 450 (TONBO biosciences) prior to 4% PFA fixation to stain dead cells. Fixed cells were permeabilized in PBS with 5% donkey serum (Sigma) and 0.3% Triton X-100 for 30 minutes at room temperature. Cells were incubated with primary antibodies in 1x PBS with 5% donkey serum and 0.1% Triton X100 overnight at 4°C. The following day, cells were washed twice in 1x PBS and unconjugated antibodies were further incubated with secondary antibodies (Alexa Fluor conjugates) for 2 hours. Antibody sources and concentrations are indicated in Supplementary Table 6. Cells were analysed using an LSR Fortessa (BD Bioscience) or FACS sorted by SH800 SONY. All data were analysed with FCS Express 6 software (BD Biosciences).

#### Generation of PDX1-eGFP positive and negative cells with minimal cytokine sets for pancreatic spheroid and hepatic organoid expansion

PDX1-eGFP reporter VFG cells passage 6 was plated at 25,000 cells/cm^2^ on polystyrene cell culture plates pre-coated with un-diluted growth factor reduced Matrigel (GRF-Matrigel; Corning) and pre-seeded with 8x10^3^ per cm^2^ irradiated DR4 mouse embryonic fibroblast (MEF). The cells were cultured in BMP4 withdrawal BMP4 media (SFD medium supplemented with FGF2 [10ng/ml], VEGF [10ng/ml], EGF [10ng/ml], 0.5 mM ascorbic acid (Wako), and 4.5x10-4 M Monothioglycerol (Sigma)) and maintained under hypoxic conditions (5% O_2_/5% CO_2_/37°C) for 5 days with medium changing every other day. For generating PDX1-eGFP positive and negative fractions, cells were further differentiated in DMEM GlutaMAX (Thermo Fisher) with 1% vol/vol B27 supplement (Life Technologies), 50ng/ml FGF2, FGF7, or FGF10 for 5 days with medium changed every day. Both BMP4 withdrawal and FGFs stimulation were performed under hypoxic conditions (5% O_2_/5% CO_2_/37°C).

The single PDX1-eGFP positive and negative cells generated from the BMP4 withdrawal and FGF10 stimulated VFG culture were sorted by FACS using a SH800 (SONY). The GFP +ve cells were expanded as pancreatic spheroid and GFP –ve cells were expanded as hepatic organoids according to the protocols described^25, 26^, except that the cultures were maintained under hypoxic conditions (5% O_2_/5% CO_2_/37°C).

#### Pancreatic differentiation

VFG cells at passages 6-8 were plated at 25,000 cells/cm^2^ on polystyrene cell culture plates pre-coated with un-diluted growth factor reduced Matrigel (GRF-Matrigel; Corning) and pre-seeded with 8000 cells per cm^2^ irradiated DR4 mouse embryonic fibroblast (MEF) in the VFG medium. Day 5 expanding VFG cells were used for pancreatic differentiations under hypoxic conditions (5% O_2_/5% CO_2_/37°C) according to protocols described as below:

For protocol adapted from Ameri et al.^24^, day 5 expanding VFG cells were treated with DMEM GlutaMAX medium (Thermo Fisher) with 1% vol/vol B27 supplement (Life Technologies) as basal media throughout the differentiation, and were supplemented with 2 µM retinoic acid (Sigma) for 3 days; then with 64 ng/ml bFGF (Peprotech) and 50 ng/ml hNOGGIN (R&D) for 3 days; and finally with 64 ng/ml bFGF (Peprotech), 50 ng/ml hNOGGIN (R&D), and 0.5 µM TBP for 3 days, with the media changed every day.

For protocol adapted from Rezania et al.^14^, day 5 expanding VFG cells were exposed to exposed to MCDB 131 basal medium throughout the differentiation, and supplemented with 1.5g/L sodium bicarbonate, 1x Glutamax, 10 mM glucose, 0.5% BSA, 0.25 mM ascorbic acid (Sigma) and 50 ng/ml FGF7 (Peprotech) for 2 days; and then with 2.5g/L sodium bicarbonate, 1x Glutamax, 10 mM glucose, 2% BSA, 0.25 mM ascorbic acid, 1:200 ITS-X (Thermo Fisher), 50 ng/ml FGF7, 1 µM RA, 0.25 µM Sant-1 (Sigma), 100 nM LDN193189 (LDN; Stemgent) and 80 nM TPB (EMD Millipore) for 2 days; and finally with 2.5g/L sodium bicarbonate, 1x Glutamax, 10 mM glucose, 2% BSA, 0.25 mM ascorbic acid, 1:200 ITS-X, 2 ng/ml FGF7, 0.1 µM retinoic acid, 0.25 µM Sant-1, 200 nM LDN and 40 nM TPB for 3 days.

For protocol adapted from Nostro et al.^12^, day 5 expanding VFG cells were fed SFD media supplemented with 50 ng/ml of hFGF10, 3 ng/ml mWnt3a and 0.75 µM dorsomorphin (Sigma) for 3 days with the medium changed every day. Then, the medium was changed to DMEM GlutaMAX medium (Thermo Fisher) with 1% vol/vol B27 (Life Technologies), 50 ng/ml hFGF10 (R&D), 50 ng/ml hNOGGIN (R&D), 50µg/ml ascorbic acid (Sigma), 2 µM retinoic acid (Sigma), with or without 0.25 µM KAAD-cyclopamine (Sigma) for one day. Finally, the medium was changed to DMEM GlutaMAX medium (Thermo Fisher) with 1% vol/vol B27 supplement (Life Technologies), 50 ng/ml hNOGGIN (R&D), 50 ng/ml hEGF (Peprotech), 10 mM nicotinamide (Sigma), and 50 µg/ml ascorbic acid (Sigma) for 4 days with the medium changed every day.

The protocol adapted from Nostro et al.^12^ was used to assess efficiency of pancreatic differentiation transient ADE cells, VFG passages 3, 6, and 12 cells generated from the PDX-eGFP reporter. The Day 5 transient ADE cells were generated as described previously and directly used for the differentiation.

### Total mRNA purification, reverse transcription and quantitative PCR Analysis

Two hundred thousand cells were washed in 1x PBS twice, lysed in RLT buffer (QIAGEN RNeasy Micro kit) containing 1% β-mercaptoethanol (Sigma) and stored at −80°C until processing. Total mRNA was isolated using the RNeasy kit according to the manufacturers’ instructions and digested with DNase I (QIAGEN) to remove genomic DNA. First strand cDNA synthesis was performed with Superscript III system (Thermo Fisher) using random primers (Thermo Fisher) and amplified using SYBR-Green (Thermo Fisher). PCR primers were designed using Primer3Plus ^61^ and validated for efficiency ranging between 95-105%. The primer sequences of the genes used in qRT-PCR are listed in Supplementary Table 6. StepOnePLUS Real-Time PCR System (Thermo Fisher) was used for qRT-PCR in 96 well plates format. Expression values for each gene were normalized against *ACTB*, using the delta-delta CT method.

### Sample preparation for Bulk RNA-seq

Total mRNA amount and RNA integrity were assessed using a Fragment Analyzer (AATI). Ribosomal RNA was removed from the samples using the NEBNext Poly(A) mRNA Magnetic Isolation Module (NEB). Sequencing libraries were prepared from 100ng of purified total mRNA using NEBNext Ultra II RNA Library Prep Kit for Illumina (NEB) according to the manufacturer’s instructions. RNA-seq libraries were sequenced for 75 cycles in single-end mode on NextSeq 500 platform (Illumina, FC-404-2005).

### Sample preparation for ATAC-seq

Dissociated single cells were washed with ice-cold PBS and pelleted at 500 x g for 10 minutes at 4°C. Fifty thousand cells were taken from a diluted stock in PBS buffer to prepare ATAC-seq libraries as described in Buenrostro et al.^62^ with slight modifications. Nuclei were prepared by resuspending the cells in 100ul ice-cold ATAC lysis buffer (10 mM Tris-HCl pH 7.4, 10mM NaCl, 3 mM MgCl2 and 0.1 % NP40) followed by incubation on ice for 15 minutes while mixing every 5 minutes. Nuclei were then collected by centrifuging at 1000 x g for 10 minutes at 4°C and pellet was then resuspended in 50 µl transposition buffer (10 mM Tris pH 8, 5 mM MgCl2 and 10% dimethylformamide). Tagmentation was performed by adding 2.5 µl Tn5 transposase (Illumina, cat #20034197) to the tubes and incubating at 37°C while shaking at 1000 rpm. Tagmentation reactions were stopped and purified with MinElute PCR Purification Kit (Qiagen, cat #28004) and tagmented DNA was eluted in10uL elution buffer (10 mM Tris pH 8.0). A PCR reaction of 50ul was assembled containing 10 µl of tagmented DNA, 2 µl NEB-Next High-Fidelity PCR Mix (NEB, cat # *M0541S)*, 5 µl of SYBR Green (Invitrogen™, cat #S7563, 50X working stock) and primers at 2 µM concentration. 10 µls of each PCR reaction was used to decide the optimum number of PCR cycles required with following conditions: 5 minutes at 72C; 30 seconds at 98°C; and 20 cycles of 10 sec at 98°C, 30 sec at 63°C and 60 sec at 72°C. The reaction was monitored in LightCycler-480 qPCR (Roche) and number of cycles required was deduced from the amplification curve. The remaining PCR reaction was then subjected to the number of PCR cycles decided above. Finally, the PCR reactions were purified with an equal volume of AMPure XP beads (Beckman, cat #A63880) following manufacturer’s protocol and eluted in 20 µl Tris pH 7.8. Libraries were quantified with Qbit dsDNA High-fidelity Assay (Invitrogen, cat # Q32851) and fragment profiles was checked using Bioanalyzer High Sensitivity assay (Agilent). Samples that showed nucleosomal bands were sequenced for 75-150 cycles in paired-end mode on an Illumina HiSeq-2000 platform.

### Generation of shRNA knockdown VFG cell lines

Short hairpin (shRNA) targeting *HHEX*, *FOXA1*, and *FOXA2* transcripts were designed using RNAi consortium (TRC) GPP Web Portal (Broad Institute) (https://portals.broadinstitute.org/gpp/public) (*HHEX*, *FOXA1*, and *FOXA2* shRNA sequences, see Supplementary Table 6). A vector delivering a scrambled sequence was used as control (scrambled shRNA sequence, see Supplementary Table 6). All shRNA sequences were cloned into a lentiviral vector (pL-U6-sgRNA-SFFV-Puro-P2A-EGFP), a gift from Kristian Helin (Addgene #12247)^63^, using *BsmBI* sites. HEK293FT packaging cells were co-transfected with the pL-U6-sgRNA-SFFV-Puro-P2A-EGFP carrying individual shRNAs and pAX8 and pCMV-VSV using lipofectamine 2000 supplemented with PEI (polyethyleneimine) according to standard protocols. SFD medium carrying lentivirus produced from HEK293FT cells (48 hrs post-transduction) was applied 1:1 with fresh VFG expansion media to one 12-well of day 2 VFG cell culture (passaged at 25,000 cells/cm^2^ at day 0). Transduction was performed in presence of polybrene at 8 µg/ml. Forty-eight hours after transduction with the sgRNA-encoding lentiviral plasmids, the VFG cells were selected and maintained at 0.25 µg/ml puromycin in standard VFG condition.

### ChIP-qPCR

Chromatin immunoprecipitation (ChIP) was carried out using the True MicroChIP kit (Diagenode) with modifications. One hundred thousand of CD184-CD117 double-positive cells sorted in the scrambled shRNA control, FOXA1, and FOXA2 shRNA knockdowns VFG culture was fixed in 1% formaldehyde (Thermo Scientific, 28906) in VFG media for 10 minutes at room temperature followed by a 5-minute quench with glycine (in True MicroChIP kit, Diagenode) at room temperature. The cells were lysed and immunoprecipitation were performed using the True MicroChIP kit (Diagenode, AB-002-0016) with the following modifications. Up to 100,000 cells were sonicated in one lysate and split into 50,000 equivalents after sonication. Samples were lysed using 50 µl of buffer tL1 and incubated for 5 minutes on ice. One hundred fifty microliters of Hank’s buffered salt solution with 1x protease inhibitor cocktail (in True MicroChIP kit, Diagenode) was added, and the lysate was sonicated in 0.65 ml Bioruptor Pico Microtubes (Diagenode). Chromatin was sheared using a Bioruptor Pico (Diagenode) with 10 cycles (30 sec on, 30 sec off). Sonicate was aliquoted in 100 µL (for 50,000 cells), and an equivalent volume of complete ChIP buffer tC1 was added. For immunoprecipitation, the following antibodies and amounts of antibody were used for the 50,000-cell ChIP: 2 µg of FOXA1 (Abcam, ab170933), 2 µg of H3K4me1 (Abcam, ab8895), and 2 µg of H3K27ac (Abcam, ab4279). Immunoprecipitation and washes were done as described in the True MicroChIP protocol, with purification using phenol chloroform extraction and ethanol precipitation. The pull-down DNA was eluted in 100µl elution buffer and qPCR was performed as described in the True MicroChIP protocol for different genomic loci. Enrichment was calculated as percentage of input. The primer sequences of the genes used in ChIP-PCR are listed in Supplementary Table 6.

### *in vitro* scRNA-seq analysis

Sequences were mapped to human hg38 genome, de-multiplexed and filtered as previously described^60, 64^ extracting a set of UMIs that define distinct transcripts in single cells for further processing. We estimated the level of spurious UMIs in the data using statistics on empty MARS-seq wells as previously described^65^. Mapping of reads was done using HISAT (version 0.1.6)^66^. Reads with multiple mapping positions were excluded. Reads were associated with genes if they mapped to an exon, using the hg38 reference human genome. The raw counts were further analyzed using Seurat (4.0.1)^67^ (https://satijalab.org/seurat/). Cells were filtered with the following thresholds (lower bound: 2,000 UMIs; 550 genes and upper bound: 35,000 UMIs; 4,950 genes). Additionally, cells with more than 20% of mitochondria content were removed as well. In Extended Fig. 1a, we subset ADE and VFG cells (505 cells). Raw counts are further normalized, log-transformed and scaled using *NormalizeData* and *ScaleData* respectively. PCA was computed on 2,000 highly variable genes without cell cycle regression. The dataset was clustered using Louvain with 0.7 resolution followed by UMAP dimension reduction on top 20 PCs. In Extended Fig. 1i, we subset for treated and withdrawal cells (562 cells). We follow the same steps above adjusting only clustering resolution set to 0.5. Detailed analyses can be found in https://github.com/brickmanlab/wong_et_al_2022/.

### *In vivo* scRNA-seq re-analysis

Li et al.^21^ dataset HRA000280 was downloaded from Genome Sequence Archive. Cells with low quality and mitochondrial content higher than 20% were filtered out (lower bound: 3,000 genes and upper bound: 9,000 genes; 400,000 UMIs). Additionally, cells labelled as “Poor quality“ were also discarded. We follow same preprocessing steps as mentioned above without clustering. We subset the final dataset for hMG, hHG, hFG and hAL population.

### Cluster alignment tool (CAT)

We use CAT to determine similarity between clusters from *in vivo* and *in vitro* studies. In short, the tool calculates mean gene expression of randomly sampled cells with replacement for each cluster 1,000 times. Euclidian distance is measured between all pairs of clusters. Small distance represents high similarity. Detailed explanation of the method can be found in Rothova et al. (submitted).

### Analysis of bulk RNA-seq data

Fastq files from bulk RNA-seq samples were aligned to the hg38/GRCh38 genome using STAR v2.5.3a^68^. Transcript expression levels were estimated with the -- quantMode GeneCounts option and GRCh38p10.v27 annotations. FastQC v0.11.7 (http://www.bioinformatics.babraham.ac.uk/projects/fastqc) was used for QC metrics, and multiqc v1.7^69^ for reporting. Data analysis was then performed with R/Biocondutcor^70^ (https://www.R-project.org). Normalization was performed with DEseq2 v1.24.0^71^. Lee et al.^34^ dataset was retrieved from NCBI GEO (GSE114102) and analyzed as above. Differential Gene Expression was performed using DESEq2 R package version 1.32.0. Z-scoring was calculated as previously described for each dataset separately.

### Processing of ATAC-seq datasets

Quality of the reads was assessed with FastQC (https://www.bioinformatics.babraham.ac.uk/projects/fastqc/) followed by trimming of poor-quality base calls and adaptor sequences with cutadapt^71^. Read-pairs were then aligned to the hg19 reference genome using bowtie2^72^ with the following parameters: *bowtie2 --no-discordant --no-mixed --no-unal --very-sensitive -X 2000.* Samtools^73^ was used for sorting alignments and format conversions. Alignments from PCR duplicates were removed using Picard (http://broadinstitute.github.io/picard/). Alignments were then converted into BED format using bedtools^74^. The 5’ ends of the reads were offset by +4 bases for the reads on Watson strand and by -5 bases for the reads on Crick strand, to reflect the exact location of Tn5 insertion site. Single-base genome-wide coverage was computed using a 30 bp fragment centred at the Tn5 insertion site in BigWig format. We called peaks using Macs2^75^ with following parameters: *macs2 callpeak --nomodel --extsize 150 --shift -75 -g ‘hs’ -p 0.01*. For each condition, data from two biological replicates was used to create a set of highly reproducible peaks Irreproducible Discovery Rate (IDR<= 0.05, ref^76^). Deeptools^77^ was employed to compute Pearson’s correlation among the conditions/replicates and for PCA plots. Bedtools *intersect* command was used to find overlapping or unique (with parameter “-v”) enhancer positions (bed format) between two conditions in question (Fig. 4a).

### Detection of differential chromatin accessibility and temporal dynamics of enhancers from ATAC-seq data

A consensus set of ATAC-seq peaks was created using reproducible peaks from all five stages of differentiation. Next, we computed normalized read coverage (rpkm) for the consensus peak-set in all stages. General Linear Modelling (GLM) was applied to the normalized counts from the step above in order to detect changes in chromatin accessibility across the stages and in both directions. We used the following parameters for differential accessibility: log2-fold > 2 or log2-fold < -2 at adjusted p-value < 0.005 (TC-seq, ref^29^). We then defined stage-specific peaks using c-means clustering of dynamic peak-set from the step above. We called 8 clusters that gave a functionally relevant pattern along the timeline of differentiation. Some clusters were merged, as they were too similar to be dealt with separately. This led to formation of the six groups of dynamic enhancers (Fig. 3c, right). RPKM normalized BigWig tracks from merged replicates were used to plot heatmaps in deeptools^77^. For locus-specific visualizations, we used UCSC Genome Browser (http://genome.ucsc.edu, ref^78^) to load BigWig tracks.

### Enrichment scoring of defined ATAC-clusters from the mapped gene sets that are up or down regulated at the PE stage compared to VFGp6

ATAC-seq peaks were assigned to genes using GREAT^79^ with the setting of single nearest gene within 25kb or 200kb (Supplementary Table 1a). The enrichment of gene-annotated ATAC-clusters in differential expression gene sets was calculated by Log 2 ratio between number of observed overlapped genes and number of expected overlapped genes from the dataset. We compared the impact of very low levels of background gene expression noise (those genes not reaching greater than 100 or 1000 reads in a particular sample, baseMean 100 or 1000) on these gene sets (Supplementary Table 2, c-f). While filtering out gene expression noise reduces the size of the gene set, it can be expanded by considering enhancers located within 200 Kb of a target gene.

### Motif analysis from ATAC-clusters

Enrichment of known and de-novo transcription factor binding motifs was calculated with the HOMER v4.11.1 suite^80^ using the *findMotifsGenome* function with default parameter.

### Hierarchical K-means clustering of expression patterns of genes annotated to ATAC-peaks clusters

Bulk RNA-seq gene expression levels were normalized using DESEq2 R package version 1.32.0^71^. The mean of normalized expression was calculated for each condition and transformed into z-scores. Gene expression levels were then separated into the different annotated ATAC-peaks clusters. Finally, gene expression patterns were grouped using hierarchical clustering (k = 10) based on Euclidian distances.

### Mapping and analysis of H3K27ac data from human embryo samples

Preprocessing and alignment of ChIP-seq reads was as described in Gerrard et al.^37^. In this study, single-end reads were aligned to hg19 genome assembly with bowtie 1.0.0 (parameters: -m1 –n 2 –l 28, uniquely mapped reads only). These alignments were received in compressed BAM format from European Genome-Phenome Archive (https://ega-archive.org/) under accession no: EGAS00001003163. We converted the alignments in to BED format and called peaks with HOMER (parameters: findPeaks - style histone) against a pooled input sample. We then used bedtools-2.30^74^ to select the peaks present in both replicates (bedtools intersect –f 0.50 –r –u –wa -a rep1.bed –b rep2.bed).

To construct a dataset for the lineage specific H3K27ac, we concatenated specific tissues peaks as follows: ectoderm (RPE; BRAIN), endoderm (PANC; LIVER; LUNG) and mesoderm (HEART; ADR). To identify unique regions of each gem layer, we use bedtools with option intersect, followed by sorting the regions using sort option and finally merging smaller regions which are subset of larger regions using option merge. This setup ensures unique count of peaks even if it’s part of a larger region. Finally, we construct files with only unique regions for each tissue again using bedtools^74^ intersect. To map different ATAC clusters to the H3K27ac datasets described above, we break *in vitro* peak files based on their class. This information is stored in the last column of the file. We then count shared peaks between *in vitro* and *in vivo* group using bedtools with the intersect option.

#### Enrichment scoring of dynamic ATAC-seq clusters with H3K27Ac regions from human embryonic tissues

The enrichment of ATAC-clusters in different lineage- and tissue-specific H3K27ac groups was calculated based on the ratio between number of observed overlapped regions (between ATAC and H3K27ac peaks) and number of expected overlapped regions from the datasets.

### Statistical Analysis

Data representation and statistical analyses were performed using GraphPad Prism. Unless mentioned otherwise, data are shown as mean ± SEM and *N* numbers refer to biologically independent replicates. Statistical significance (*P*<0.05) was determined as indicated in figure legends using one-way ANOVA Tukey’s multiple comparison test (Figs. 1c and 1d), one-way ANOVA Dunnett’s multiple comparison test (Figs. 2a, 2b, 2e, and 6e; Extended Figs. 1m, 1n, and 2d), and unpaired two-tailed t-test (Figs. 2d, 6b, 6c, and 6d; Extended Figs. 1d, 6c, and 6e).

### Data and Code Availability

Sequencing data generated in this study are available on NCBI GEO with accession GSE-185670 (bulk RNA-seq), GSE188362 (single-cell RNA-seq), and GSE108623 (ATAC-seq). H3K27ac ChIP-seq dataset of human embryos was from our previous study (Gerrard et al.^37^). Code used to perform the analyses in this study is available at https://github.com/brickmanlab/wong_et_al_2022/ or from the corresponding authors upon request.

### Materials Availability

Cell lines and reagents generated for this study are available from the lead contact with a complete Materials Transfer Agreement.

### Lead Contact

Further information and requests for resources and reagents should be directed to and will be fulfilled by the lead contact, Joshua Mark Brickman.

## Acknowledgements

We thank Paul Gadue for sharing protocols for EP expansion; Henrik Semb for the HUES4 WT and PDXeG clone 170-3 cell lines; we thank the DanStem Genomics Platform, DanStem Flow Cytometry Platform, the DanStem Imaging Platform and the DanStem Stem Cell Culture Platform for training, technical expertise, support and the use of instruments. We also thank members of the Brickman and Bickmore labs for critical comments on this manuscript. We are grateful to Anne G. Botton for critical reading of the manuscript. This work was funded by HumEn under the European Union Seventh Framework Programme FP7/2007-2013 (HEALTH-F4-2013-602889) and by the Danmarks Frie Forskningsfond (DFF-6110-00009). W.A.B is supported by MRC University Unit grant (MC_UU_00007/2). N.A.H. is supported by MRC (MR/000638/1 and MR/S036121/1). R.E.J. is supported by Diabetes UK Harry Keen Clinician Scientist fellow. J.A.R.H is supported by the Novo Nordisk Foundation (grant number NNF20OC0063268). The Novo Nordisk Foundation Center for Stem Cell Biology was supported by the Novo Nordisk Foundation (grant number NNF17CC0027852) and the Novo Nordisk Foundation Center for Stem Cell Medicine is supported by Novo Nordisk Foundation (grant number NNF21CC0073729).

## Author contributions

Y.F.W., W.A.B., and J.M.B. conceived of the project. Y.F.W. and Y.K. conducted experiments. W.A.B. and J.M.B. designed the experiments and obtained funding for the study. Y.F.W., Y.K., M.P., J.A.R.H., M.M.R., R.S.M., and S.P. performed data analysis. M.M.R. conducted the single-cell sequencing experiment. N.A.H. and R.E.J. provided insight into organ specific enhancer regulation and H3K27ac ChIP-seq data from human embryos. Y.F.W., Y.K., W.A.B., and J.M.B. wrote the manuscript.

---

**Extended Data Fig. 1.**
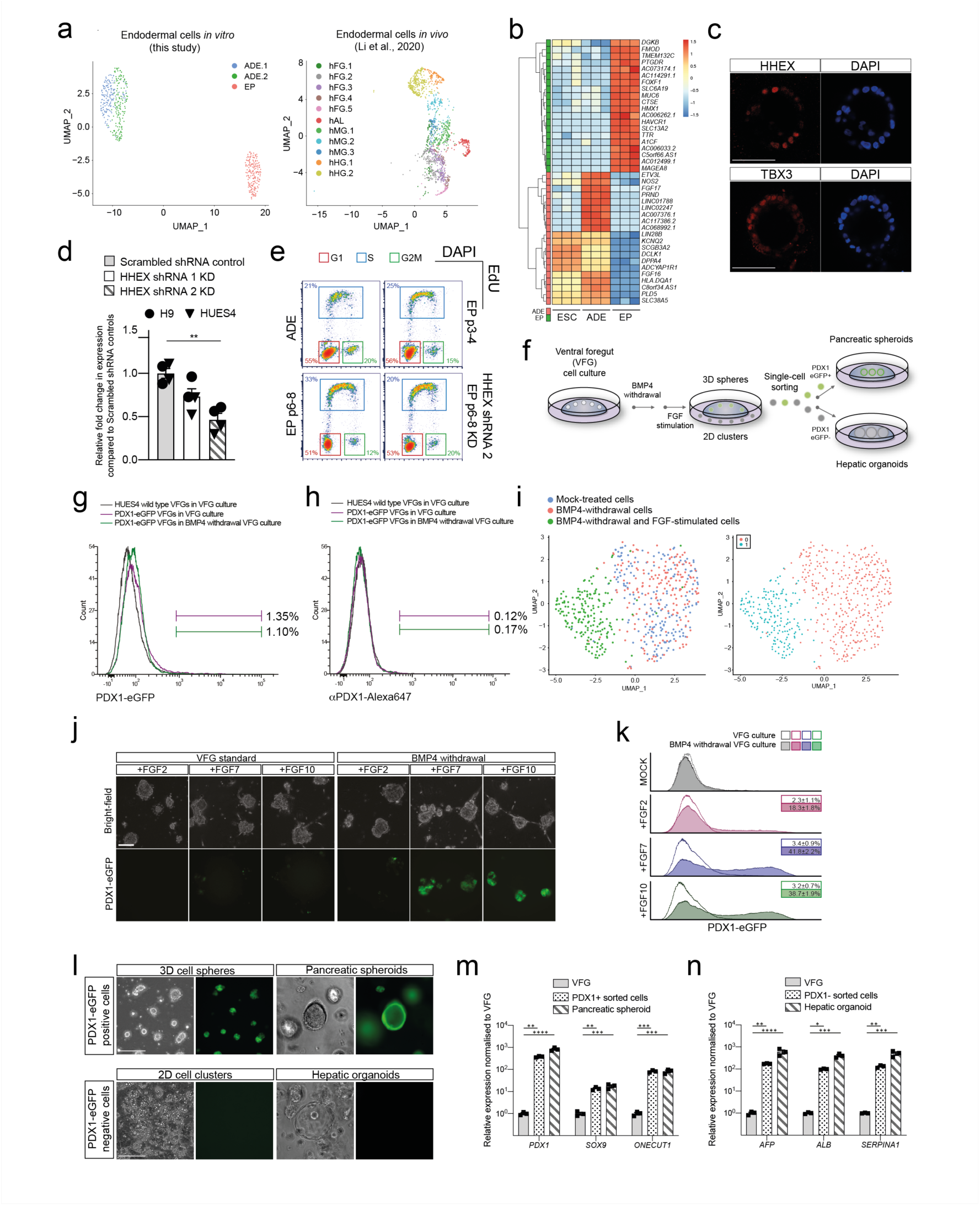
Ventral foregut identity of expanding endodermal progenitors. **a**, Left: UMAP visualization of single cells from the transient ADE (ADE.1 and ADE.2) and EP passage 6 samples. Right: UMAP visualization of single cells from different endodermal populations from early human embryos reported in Li et al.^21^. **b**, Heatmap illustrating gene expression in H9-derived ESC, ADE, and EP cells (*N*=3) from bulk RNA-seq dataset. Log normalized counts of the top 20 differentially expressed genes for each condition (ADE vs EP) are shown. **c**, Representative immunostaining of hAL markers, HHEX and TBX3, in EP passage 6 cells derived from H9 ESC cells. **d**, Expression analysis in HHEX shRNA KD cells (set 1 and set 2) by qRT-PCR. Relative fold change in mRNA of HHEX gene in KDs and control VFG cells was assayed by qRT-PCR. Expression is normalized to *ACTB*. Data are represented as mean ± SEM (*N*=4). Statistics analysis (**P<0.01, unpaired two-tailed t-test) was performed between KD and control VFG cells. Comparisons without an indicated P value are not significant. **e**, Representative flow cytometry plots used to analyse the cell cycle in transient ADE, early EP/VFG (p3-4), expanding EP/VFG (p6-8) and HHEX depleted EP/VFG (p6-8). Cells were stained with EdU and DAPI. Cells in G1 (red), S (blue), and G2M (green) were subgated and percentages of each fraction shown. **f**, Schematic representation showing the conversion of VFG culture to pancreatic and hepatic expansion. The figure illustrates the generation of PDX1-eGFP positive (PDX1+) and negative (PDX1-) cells from VFG culture after BMP4 withdrawal and subsequent stimulation with FGF. **g**, Flow cytometry of eGFP expression for HUES4 wild type VFGs (grey), and the PDX1-eGFP reporter VFGs (purple) in VFG culture, and the PDX1-eGFP reporter following BMP4 withdrawal (green). Fractions of PDX1+ve were gated and percentages are shown. **h**, Intracellular flow cytometry for PDX1 (with Alexa647 conjugation) in HUES4 wild type VFGs (grey), PDX1-eGFP reporter VFGs (purple), and following BMP4 withdrawal from VFG culture for the PDX1-eGFP reporter (green). Fractions of PDX1+ve were gated and percentages were shown. **i**, Left: UMAP visualisation of 526 cells isolated from mock-treated VFG (blue), VFG cells grown in the absence of BMP4 (red), and transient pancreatic induction by FGF2 simulation (green). Right: UMAP visualisation of Seurat clustering from the samples described on the left. **j**, Representative bright- field (top) and fluorescent (bottom) images for the PDX1-eGFP reporter VFGs cultured in standard VFG (left) or BMP4 withdrawal (right), and then treated with FGF2, FGF7, or FGF10. Scale bar = 50 µm. **k**, Flow cytometry of eGFP expression for the conditions described in (**g**), including mock-treated cells. Percentages of PDX1+ve cells were shown in the rectangle boxes of each histogram. **l**, Top row: PDX1+ cells form 3D spheres and then expand as pancreatic spheroids. Scale bar = 50µm. Bottom row: PDX1-cells form 2D clusters and then expand as hepatic organoids in 3D. Scale bar = 50µm. **m**, Relative fold change in mRNA of pancreatic markers (PDX1, SOX9, and ONTCUT1) in p6 VFGs, PDX1+ve cells from the differentiating 3D cell spheroids, and established pancreatic spheroid culture (as described in **k**) was assayed by qRT-PCR. Expression is normalized with *ACTB*. Data are represented as mean ± SEM (*N*=3). **P<0.01, ***P<0.001, ****P<0.0001 (one-way ANOVA Dunnett’s multiple comparison test compared with VFG). **n**, Relative fold change in mRNA of hepatic markers (AFP, ALB, and SERPINA1) in p6 VFGs, PDX1-ve cells from the differentiating 2D cell clusters, and established hepatic organoid culture (as described in **k**) was assayed by qRT-PCR. Expression is normalized to *ACTB*. Data are represented as mean ± SEM (*N*=3). *P<0.05, **P<0.01, ***P<0.001, ****P<0.001 (one-way ANOVA Dunnett’s multiple comparison test compared with VFG).

**Extended Data Fig. 2.**
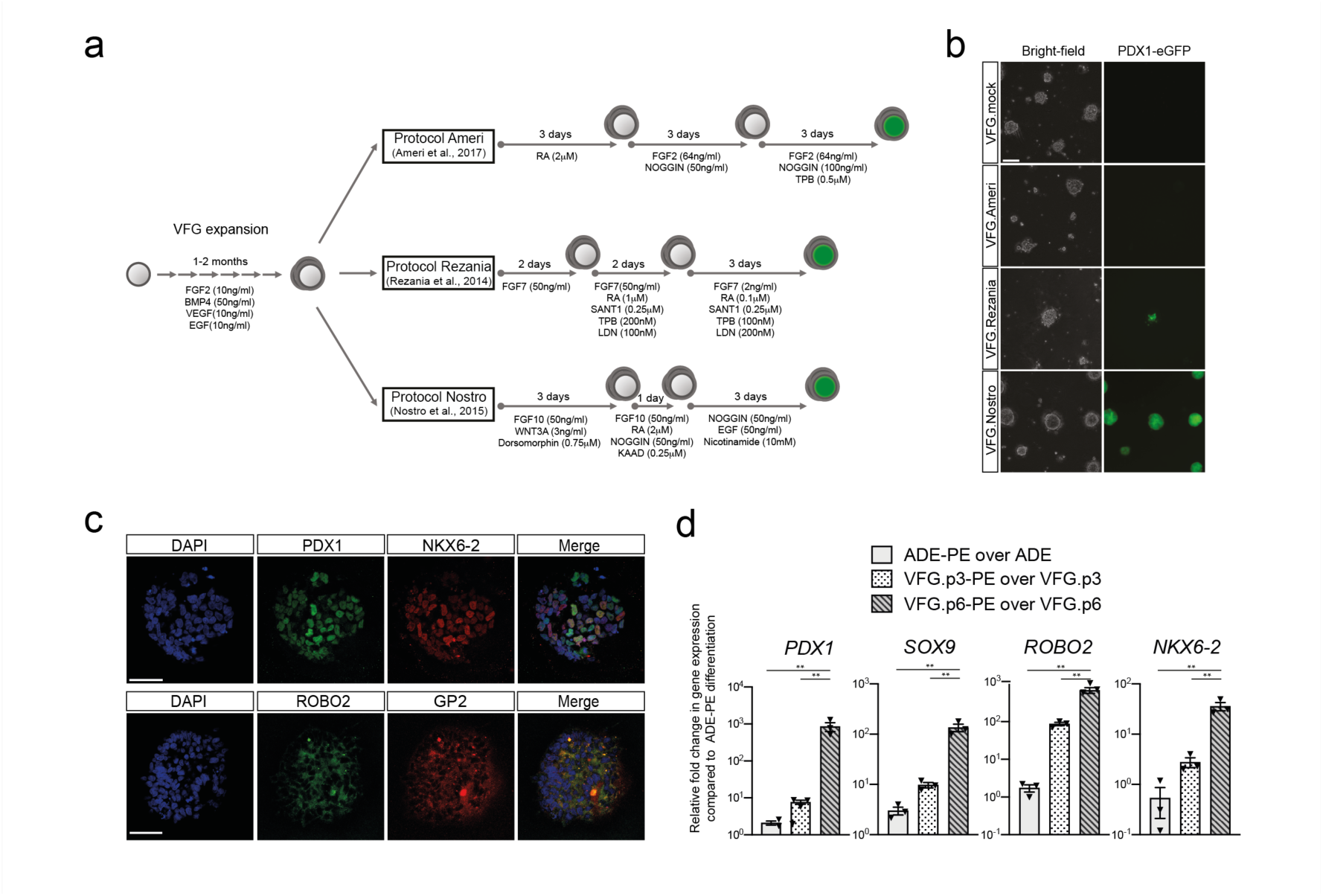
*in vitro* differentiation of VFG culture towards pancreatic endoderm. **a**, Schematic diagram for stepwise pancreatic differentiation with protocols from Ameri et al.^24^, Rezania et al.^14^ and Nostro et al.^12^ from established VFG culture. **b**, Representative bright-field (left) and fluorescent (right) images for the PDX1-eGFP reporter VFGs differentiated with protocols from Ameri et al.^24^, Rezania et al.^14^ and Nostro et al.^12^. Scale bar = 50 µm. **c**, Representative immunostaining of PDX1 (green) and NKX6-2 (red) in the top row; ROBO2 (green) GP2 (red) in the bottom row, including DAPI (blue) for p6 VFG cells differentiated with protocol from Nostro et al.^12^. Scale bar = 50 µm. **d**, Box plots showing relative fold change in mRNA of pancreatic markers PDX1, SOX9, ROBO2, and NKX6-2 in pancreatic differentiation from ADE and VFG at p3 and p6 cells. Data are represented as mean ± SEM (*N*=3). **P<0.01 (one-way ANOVA Dunnett’s multiple comparison test compared with PE differentiation from ADE).

**Extended Data Fig. 3.**
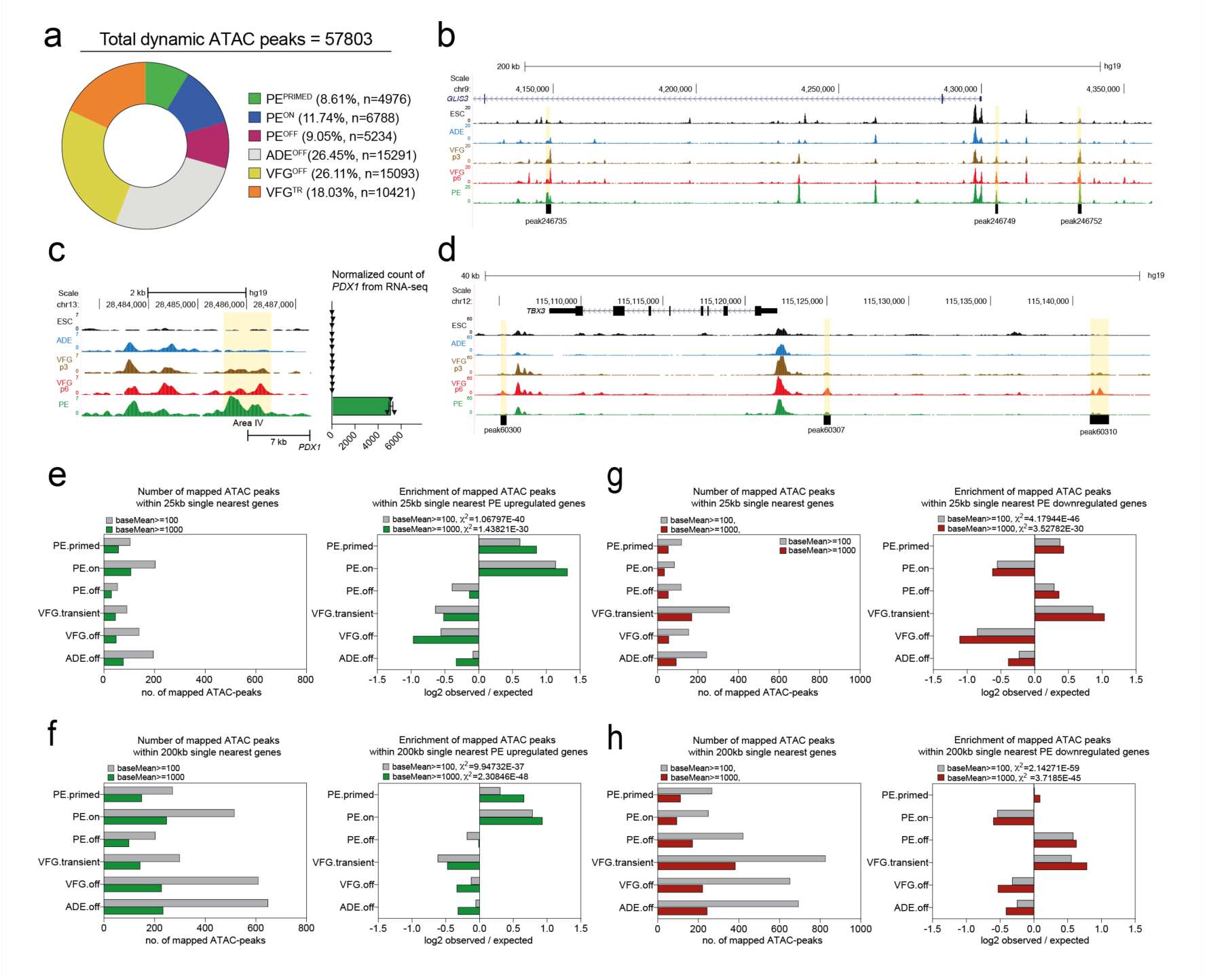
The dynamic chromatin landscape and gene expression in VFG expansion and further Differentiation. **a**, Pie-chart showing distribution of dynamic ATAC-peaks (n=57803) with percentage and numbers of peak indicated for each cluster. **b**, UCSC Genome Browser screen shot at the *GLIS3* locus showing ATAC-seq data from ESC, ADE, VFGp3, VFGp6, and PE. Genome coordinates (bp) are from the hg19 assembly of the human genome. Three PE^PRIMED^ elements (peaks 246735, 246749, and 246752) are shown at the bottom and the corresponding regions are highlighted in yellow. **c**, Left: UCSC Genome Browser screen shot at the PDX1 locus showing ATAC-seq data from ESC, ADE, VFGp3, VFGp6 and PE. Genome coordinates (bp) are from the hg19 assembly of the human genome. The region of the area IV enhancer is highlighted in yellow and the approximate distance between the region and *PDX1* TSS is indicated. Right: bar plots for normalized RNA-seq counts (normalized read count) for *PDX1* across ESC, ADE, VFGp3, VFGp6, and PE cells. **d**, UCSC Genome Browser screen shot at the *TBX3* locus showing ATAC-seq data from ESC, ADE, VFGp3, VFGp6, and PE. Genome coordinates (bp) are from the hg19 assembly of the human genome. Three VFG^TR^ elements (peaks 60300, 60307, and 60310) are shown at the bottom and the corresponding regions are highlighted in yellow. **e**, Mapping dynamic enhancer classes to gene expression (up-regulated genes). Left: Number of mapped ATAC peaks in each cluster defined in 3c located within 25 Kb of the single nearest gene’s TSS from the PE up-regulated gene set with baseMean >100 (grey) or >1000 (green). Right: Enrichment (log2 observed/expected) of the PE up-regulated gene set with baseMean > 100 (grey) or >1000 (green), in proximity (within a 25 Kb window) to ATAC-clusters defined in Fig. 3c. All data shown are significant by chi-square analysis. **f**, Mapping dynamic enhancer classes to gene expression (down-regulated genes). Left: Number of mapped ATAC peaks in each cluster defined in Fig. 3c located within 200 Kb of the single nearest gene’s TSS from the PE up-regulated gene set with baseMean >100 (grey) or >1000 (green). A complete set of data was listed in Supplementary Table 1c and d. Right: Enrichment (log2 observed/expected) of the PE up-regulated gene set with baseMean > 100 (grey) or >1000 (green), in proximity (within a 200 Kb window) to ATAC-clusters defined in Fig. 3c. All data shown are significant by chi-square analysis. **g**, Mapping dynamic enhancer classes to gene expression (up-regulated genes). Left: Number of mapped ATAC peaks in each cluster defined in Fig. 3c located within 25 Kb of the single nearest gene’s TSS from the PE down-regulated gene set with baseMean >100 (grey) or >1000 (green). Right: Enrichment (log2 observed/expected) of the PE down-regulated gene set with baseMean >100 (grey) or >1000 (green), in proximity (within a 25 Kb window) to ATAC-clusters defined in Fig. 3c. All data shown are significant by chi-square analysis. **h**, Mapping dynamic enhancer classes to gene expression (down-regulated genes). Left: Number of mapped ATAC peaks in each cluster defined in Fig. 3c located within 200 Kb of the single nearest gene’s TSS from the PE down-regulated gene set with baseMean >100 (grey) or >1000 (green). A complete set of data was listed in Supplementary Table 1, e and f. Right: Enrichment (log2 observed/expected) of the PE down-regulated gene set with baseMean > 100 (grey) or >1000 (green), in proximity (within a 200 Kb window) to ATAC-clusters defined in Fig. 3c. All data shown are significant by chi-square analysis.

**Extended Data Fig. 4.**
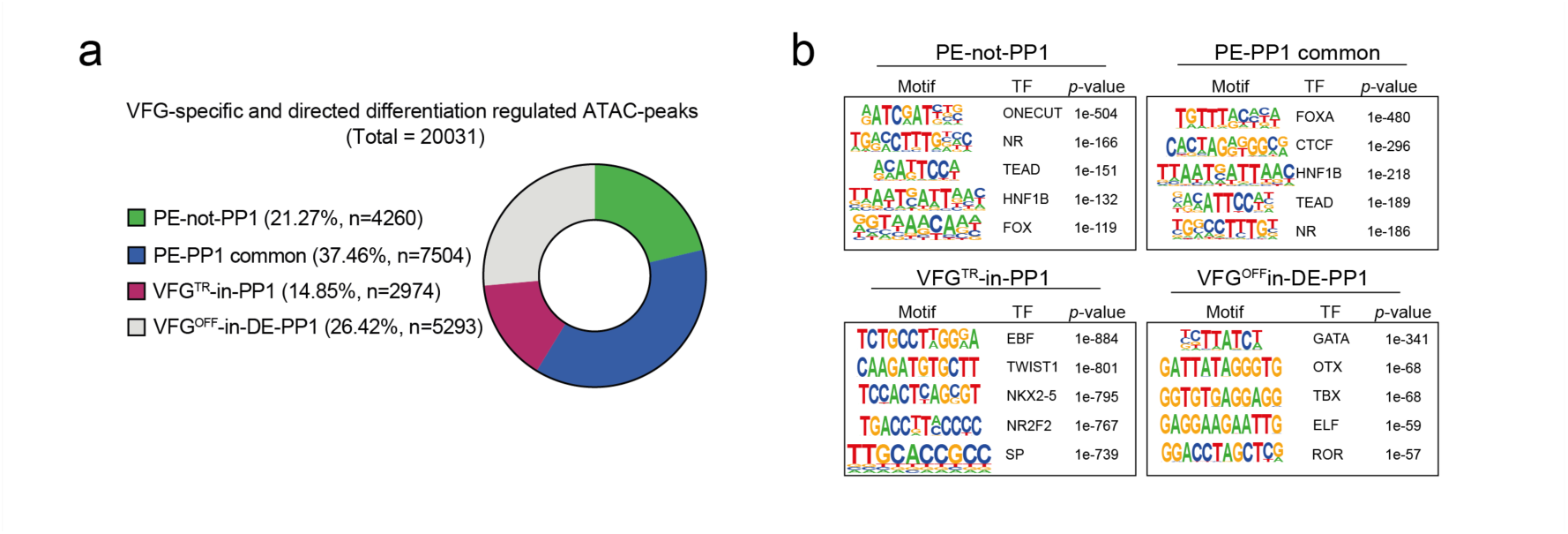
Distribution and motif analysis of VFG-specific and directed differentiation regulated ATAC-peak clusters. **a**, Pie-chart showing distribution of VFG-specific and directed differentiation regulated ATAC-peaks (n=20031) with percentage and number of peaks indicated for each cluster. **b**, HOMER motif analysis for VFG-specific and directed differentiation regulated ATAC-peak clusters. The top five de novo motifs are shown.

**Extended Data Fig. 5.**
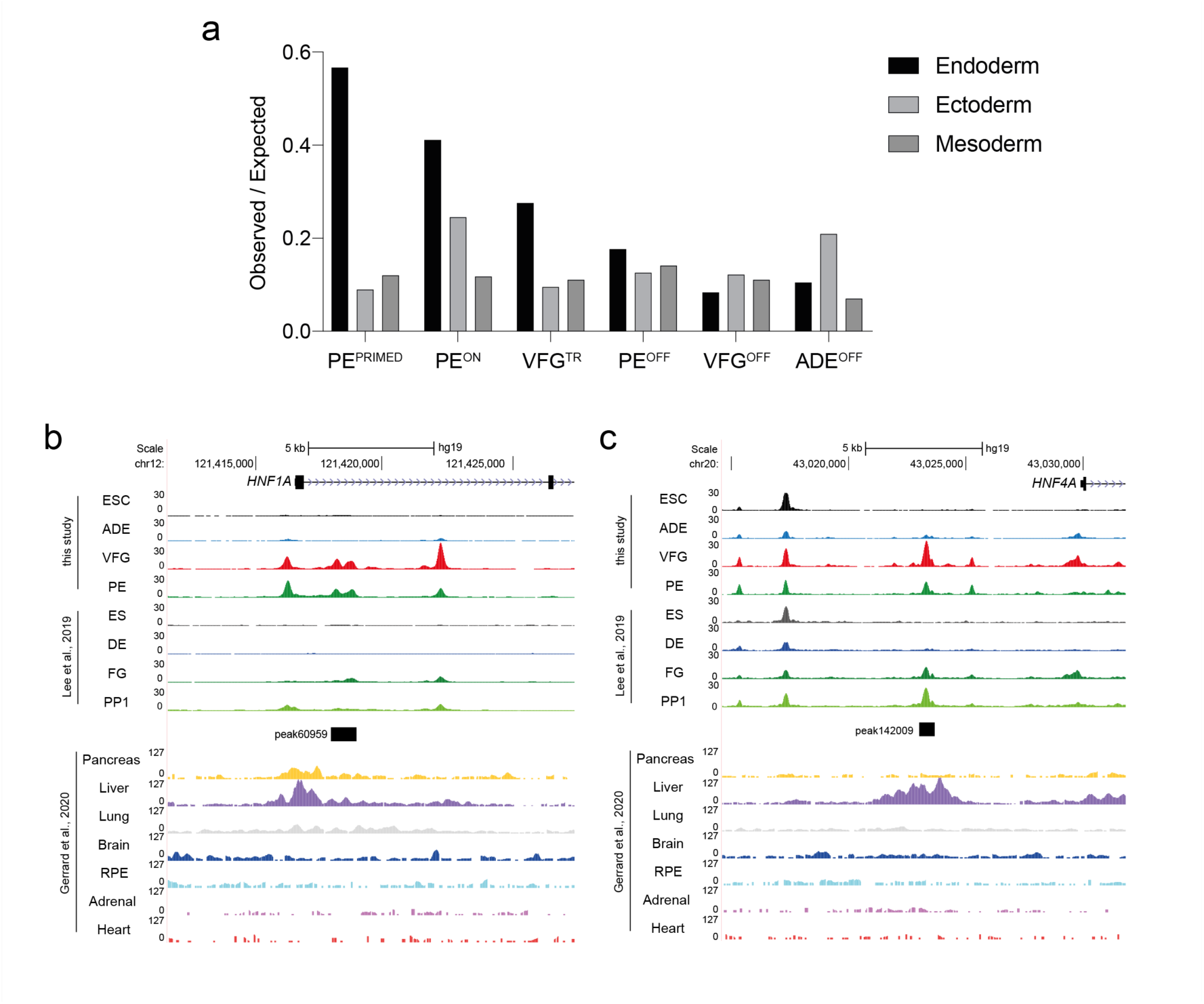
VFG expansion insures higher fidelity regulation of enhancers normally exploited in fetal organogenesis. **a**, Enrichment of lineage-specific H3K27ac enhancers (endoderm, ectoderm, and mesoderm) from human embryos^37^ observed in the different ATAC clusters defined in this study displayed by enrichment score (observed/expected) in a bar plot. **b**, UCSC Genome Browser screen shot at the *HNF1A* locus showing ATAC-seq data in this study (ESC, ADE, VFG, and PE) and from Lee et al.^34^ (ES, DE, FG, PP1) alongside H3K27ac ChIPseq data (Hanley Lab 2014, 2015, STAR_hg38_FULL, ds99 scaled)^37, 78^ from multiple human embryonic tissues (pancreas, liver, lung, brain, RPE, adrenal, and heart). Genome coordinates (bp) are from the hg19 assembly of the human genome. PE-not-PP1 element peak60959 overlaps with both pancreas- and liver-specific H3K27ac enhancers. **c**, UCSC Genome Browser screen shot at the *HNF4A* locus showing ATAC-seq data in this study (ESC, ADE, VFG, and PE), from Lee et al.^34^ (ES, DE, FG, PP1) alongside H3K27ac ChIP-seq (Hanley Lab 2014, 2015, STAR_hg38_FULL, ds99 scaled)^37, 78^ from multiple human embryonic tissues (pancreas, liver, lung, brain, RPE, adrenal, and heart). Genome coordinates (bp) are from the hg19 assembly of the human genome. VFG^TR^-in-PP1 element peak142009 overlaps with a liver-specific H3K27ac enhancer.

**Extended Data Fig. 6.**
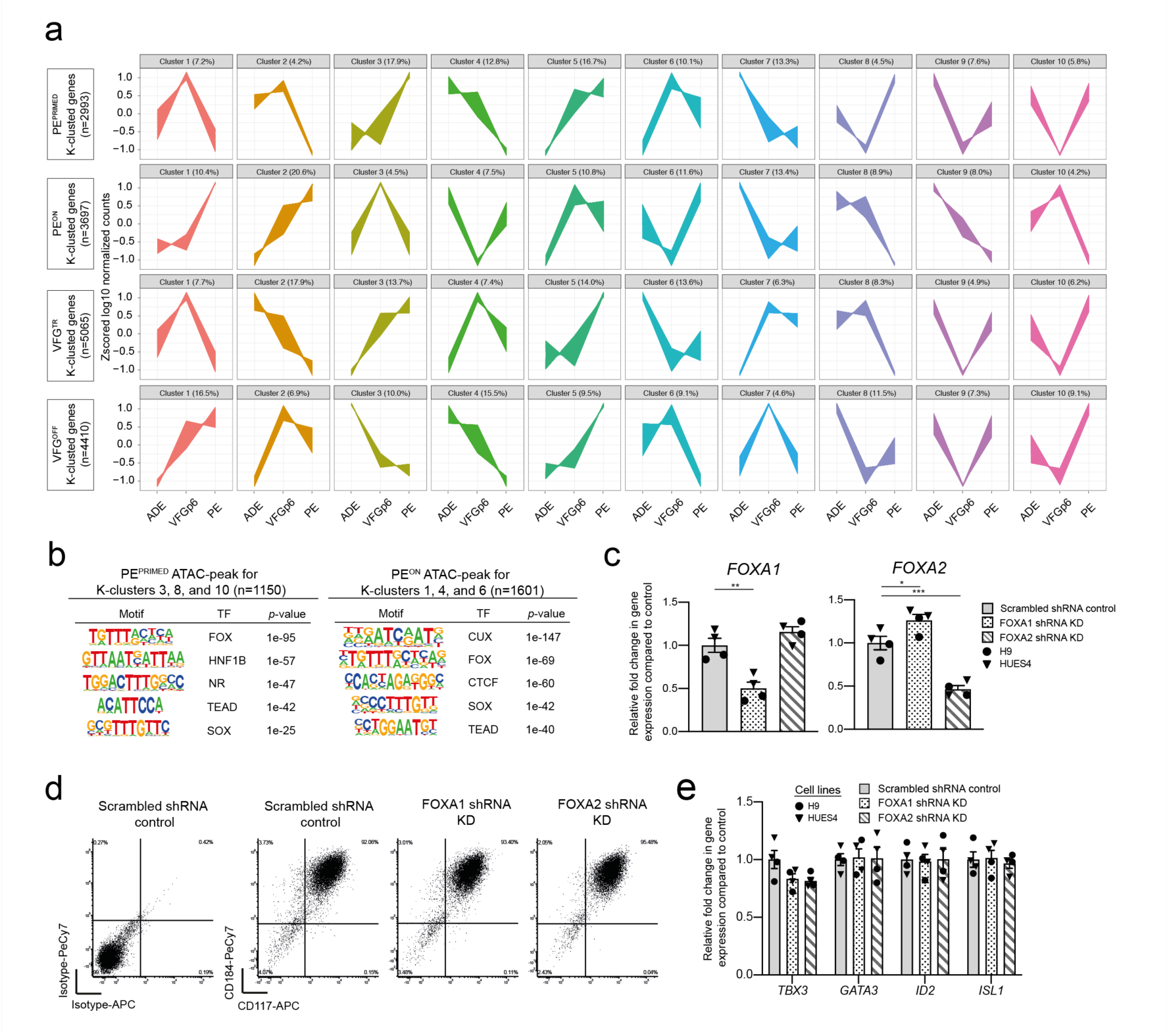
Enhancer regulation by FOXA proteins during VFG expansion and pancreatic differentiation. **a**, K-means clustering of genes within 200Kb of peaks in ATAC clusters as defined in Fig. 4a (PE^PRIMED^, PE^ON^, VFG^TR^, and VFG^OFF^). Z-scored log10 normalized gene expression of ADE, VFGp6, and PE samples were plotted for clustered genes (n=10). **b**, Homer motif analysis for genes mapped to the vicinity of PE^PRIMED^ enhancers for K-means clusters 3, 8, and 10 (left) and genes associated to PE^ON^ enhancers for K-means clusters 1, 4, and 6 (right). **c**, Expression analysis in FOXA1 and FOXA2 shRNA KD cells (described in Fig. 5B) by qRT-PCR. Expression of FOXA1 (left) and FOXA2 (right) in the KD cells was normalized relative to the expression in scrambled shRNA controls. Data are represented as mean ± SEM (*N*=4). *p<0.05, **p<0.01, ***p<0.001 (unpaired two-tailed t-test). Comparisons without an indicated p value are not significant. **d**, Representative flow cytometry density plots showing CD184 (CXCR4) and CD117 (KIT) expression in scrambled shRNA control, FOXA1, and FOXA2 shRNA KD VFG cells. Bottom left quadrant indicates gating based on isotype staining controls in scrambled shRNA control VFG cells. **e**, Expression analysis in FOXA1 and FOXA2 shRNA KD cells (described in Fig. 6b) by qRT-PCR. Expression of VFG markers TBX3, GATA3, ID2, and ISL1 in the KD cells was normalized relative to that in scrambled shRNA controls. Data are represented as mean ± SEM (*N*=4). No statistical difference (unpaired two-tailed t-test) was found in the comparisons.

## Other Supplementary Materials for this manuscript include the following

- Supplementary Table 1. Summary of dynamic ATAC peaks, and their annotated genes classified by differential expression between VFG and PE samples.
- Supplementary Table 2. Summary of VFG-specific ATAC peaks, and their annotated genes classified by differential expression between VFG and PE samples.
- Supplementary Table 3. H3K27ac datasets derived from different human embryonic tissues.
- Supplementary Table 4. Dynamic enhancers defined by ATAC-seq clusters mapped to H3K27ac datasets from human embryonic tissues.
- Supplementary Table 5. Motif enrichment results for dynamic enhancers defined by ATAC-seq clusters.
- Supplementary Table 6. Oligonucleotide sequences and antibodies used in this study.

